# Development and Characterization of a Tunable PDMS Substrate Model for Investigating Elastic Properties and Mechanical Stretching in Intervertebral Disc Cells

**DOI:** 10.1101/2025.07.07.663371

**Authors:** Johannes Hasler, Mikkael Lamoca, Kory Schimmelpfennig, Shuhuan Zhang, Wolfgang Hitzl, Sami Farajollahi, Vinay V. Abhyankar, Christopher L. Lewis, Rui Liu, Karin Wuertz-Kozak

## Abstract

**Background:** Aberrant mechanical loading and altered extracellular matrix (ECM) composition favor catabolic cell responses, contributing to intervertebral disc (IVD) degeneration and ultimately impairing the integrity of the annulus fibrosus (AF). This highlights the need for new *in vitro* models to investigate the interplay of mechanical loading and cell-substrate interactions. Therefore, this study introduces a tunable stretching chamber platform to simultaneously study both factors in AF degeneration.

**Methods:** Tunable PDMS substrates were fabricated by adjusting ratios of Sylgard 184 and Sylgard 527, enabling molding into stretching chambers or well plates. Substrates underwent mechanical, optical and chemical characterization. Bovine AF cells were seeded onto the substrates and cultured under static conditions or subjected to cyclic strain (8% strain at 1 Hz). Substrate biocompatibility and cell morphology were assessed over 72 h in static cultures. Dynamic responses were assessed by cell viability and alignment. Digital Image Correlation (DIC) was employed to assess surface strain in custom-designed and commercial (STREX) stretching chambers at strains between 8 to 20%.

**Results:** PDMS formulations resulted in a stiffness (E modulus) range of 8.72 – 238.00 kPa, demonstrating distinct viscoelastic profiles. All substrate formulations supported AF cell adhesion and viability. DIC analysis revealed non-uniform axial and transverse strain distributions on membrane surfaces. Cyclic stretching showed that cells maintained viability up to 14 h. Additionally, cells responded to the applied strain by perpendicular alignment to the stretch axis at 6 h.

**Conclusions:** The PDMS based stretching platform offers a biocompatible and tunable mechanical environment that mimics physiological and pathophysiological elastic properties. It enables systematic investigations on how elastic properties and mechanical strain modulate AF cell behavior in IVD disease progression. Finally, this study raises the awareness of non-uniform and transverse strain components within the stretching chambers and highlights the discrepancy of effective strain transfer to the cell interaction surface.

## 1 Introduction

Mechanical cues are crucial in regulating fundamental cellular functions and preserving tissue health [1, 2]. Cells continuously sense these mechanical changes through mechanosensitive ion channels and integrin dependent extracellular matrix (ECM) interaction, which transduces mechanical stimuli into biochemical signals [3–5] and thereby regulate proliferation, differentiation, and ECM remodeling. Disturbances of the ECM environment and mechanical loading can impair these cell-ECM interactions [1]. Notably, these changes have been widely implicated in various degenerative tissue pathologies, such as pulmonary [6] and liver fibrosis [7], cancer progression [8], cardiovascular diseases [9], and musculoskeletal disorders [1, 10, 11].

Mechanical loading has a significant impact on musculoskeletal tissues and regulates essential cellular functions [1]. Abnormal loading conditions, however, interfere with these functions and accelerate the disease progression [1, 12]. An important example is the intervertebral disc (IVD), which is located between the vertebral bodies, transmitting and distributing loads in the spine while being highly responsive to mechanical loading. The IVD consists of the central, gelatinous nucleus pulposus (NP), which primarily absorbs compressive loads, and exerts tensile forces to the surrounding fibrous annulus fibrosus (AF), which provides structural support through its highly aligned collagen I fibers [13]. While physiological loading of the IVD supports nutrient diffusion and induces anabolic cell responses that promote tissue health, hyper-physiological loading induces cell damage, inflammation, and catabolic responses in the IVD [14, 15]. These detrimental factors accelerate degeneration and promote structural changes that contribute to AF stability and can ultimately result in IVD herniation — a major cause of lower back pain (LBP) [13, 16]. Aside from mechanical loading, numerous studies reported that degeneration is closely linked to alterations in the mechanical properties of the AF, particularly an increase in AF stiffness [17–19], enhancing its susceptibility to rupture. Given the connection of mechanical loading and ECM stiffening in IVD degeneration [20], it is crucial to understand the molecular pathways through which mechanical cues — whether arising from ECM changes, mechanical loading or a combination of both — influence degeneration. This underlines the need for advanced tools and model systems that enables studying cellular responses to mechanical loading under physiological and pathophysiological stiffness conditions, and their role in disease progression [21].

Mechanical stretching is a prevalent factor in the AF and should be accurately replicated in *in vitro* experimental designs to reflect physiological conditions. Typically, mechanical stretching is studied using either custom-manufactured or commercially available stretching bioreactors from companies such as Flexcell or STREX [5]. These systems employ polydimethylsiloxane (PDMS) as their flexible membranes that allow elastic deformation under applied force, transmitting the mechanical stimuli to cells. Both bioreactor types are well suited for applications such as gene expression studies or live-cell imaging, [5, 22, 23] however, their systems are costly and limited information on the system characterization is available. In this study we primarily focused on the uniaxial stretching bioreactor from STREX due to availability, experimental needs and limitations of the device. The system incorporates PDMS based membranes into stretching chambers, where these membranes are seeded with the desired cell type and affixed to pins. The device then stretches the membrane at specified magnitudes and frequencies, thereby mechanically stimulating the attached cells [5, 22]. However, the system has two significant limitations. First, the chambers do not accurately mimic the physiological or pathophysiological stiffness of most tissues. Second, there is insufficient data to confirm whether cells in different areas of the chamber experience uniform strain. In comparison, Flexcell offers PDMS membranes with stiffness range between 1 to 60 kPa (CellSoft® 6-Well BioFlex®), covering a physiologically relevant stiffness range. While this is advantageous, these commercial membranes are costly, and their proprietary formulations are undisclosed. Additionally, strain distribution of the Flexcell membranes in literature has revealed non-uniform strains [24] as well as discrepancies between programmed and actual strain [25]. Most importantly, the variation of the strain magnitude and distribution on the different membrane stiffnesses remain unexplored.

PDMS is an excellent choice for biological applications and stretching devices not only due to its excellent biocompatibility, optical transparency, modifiable surface chemistry, and capacity to undergo large deformations. The mechanical properties including stiffness, can be precisely and easily modified over a very large range, and is a well characterized material that makes it accessible and simple to use, for less experienced researchers while being cost-effective [26]. Beyond that PDMS offers a broad engineering flexibility enabling the incorporation of features such as micro- or nano-scale topographies [27], stiffness gradients [28] or pores [29]. Additionally, PDMS can be easily molded into unique geometries and formats, like well plates, allowing multimodal readouts for gene expression analysis, imaging, fluorescent staining, and mechanical stimulation. This demonstrates the benefits of PDMS, which often outperforms alternative materials in terms of versatility and customization, making it a strong candidate for mechanobiological research. Despite these strengths, STREX systems do not fully capitalize on these advantages, thereby limiting their ability to replicate the nuanced mechanical environments found in physiological or pathophysiological tissues. Historically, the Sylgard 184 formulation, with its traditional 10:1 base-to-crosslinker ratio, was widely used for PDMS applications because of its well-characterized mechanical properties. However, this formulation falls short in replicating the physiological elastic properties observed in tissues such as the IVD [30]. Researchers initially attempted to widen the stiffness range by tweaking the base-to-crosslinker ratio of Sylgard 184, but this approach often led to the formation of uncrosslinked polymer sites [31]. More recently, advancements have led to the development of hybrid formulations that combine Sylgard 184 with Sylgard 527 [32]. These innovative mixtures can achieve moduli as low as 5 kPa, thereby offering a broader stiffness range [32, 33]. Importantly, they maintain key surface characteristics, including roughness, energy, protein adsorption, and wettability, without compromise [30, 32, 34].

Incorporating more advanced PDMS formulations into bioreactor systems is therefore a promising tool for mechanobiological research by enabling controlled studies of how cells react to substrate stiffness and mechanical loading. Given the significance of mechanical loading and ECM properties on cell responses, stretching platforms that combine tunable PDMS formulations with mechanical stimulation are valuable for investigating these interactions. These stiffness-adjusted PDMS stretching chambers not only allow studying the mechanical mechanisms underlying IVD health and disease but can also be adjusted to research in various other pathologies. However, despite its importance, limited studies explore the synergy of mechanical loading and substrate elastic properties [35].

This study aims to characterize the commercially available STREX uniaxial stretching system and to introduce a novel, practical, and accessible cost-effective alternative platform using non-proprietary Sylgard 527 and Sylgard 184 elastomer blends. This approach produces stretching chambers with elastic properties that spans a physiologically relevant range for the IVD. This tool enables systematic investigations on how elastic properties and *in vitro* mechanical strain modulates AF cell behavior in IVD mechanobiology. Therefore, the study hypothesized that custom PDMS substrates using Sylgard 184 and 527 better replicate physiological elastic properties than commercial stretching chambers by STREX, and that uniaxial loading of PDMS membranes influence surface strain distribution and non-uniform strains, subjecting cells to varying strains depending on their location within the chamber.

## 2 Methods

### 2.1 Fabrication of static PDMS substrates

Two commercially available silicone formulations (Sylgard 184 Silicone Elastomer Kit, Dow, 2646340 and Sylgard 527 Dielectric Gel Kit, Dow, 1696742) were used to prepare PDMS substrates with tailored stiffness (Low, Mid, High). Both formulations: Sylgard 184 and 527 were prepared according to manufacturers’ recommendations (1:10 wt ratio for Sylgard 184 and 1:1 wt ratio for Sylgard 527). As shown in Table 1, Sylgard 184 was added to Sylgard 527 at different mass ratios to modulate the substrate stiffness. Additionally, a control sample consisting of Sylgard 184 (commercial formulation) was prepared. The mixtures were mixed for 1 min, followed by a degassing step in a vacuum chamber. The PDMS mixtures were then incorporated into tissue culture plastic (TCP) plates and cured in an oven (Thermo Scientific, HERA THERM Oven) at 65°C for 24 h to ensure complete crosslinking.

**Table 1:** Representation of the blend ratios by weight (*wt.*) of Sylgard 184 and 527 formulations resulted in the PDMS formulations Low, Mid, High, Commercial.

| PDMS Formulation<br>(Stiffness) | Sylgard 184 (wt) | Sylgard 527 (wt) |
| --- | --- | --- |
| *Low* | 0% (wt.) | 100% (wt.) |
| *Mid* | 14% (wt.) | 86% (wt.) |
| *High* | 24% (wt.) | 76% (wt.) |
| *Commercial* | 100% (wt.) | 0% (wt.) |

### 2.2 Stretching chamber manufacturing

An aluminum mold (Figure 1A) was fabricated to cast PDMS stretching chambers compatible with the automated cell stretching system STB-1400 (STREX). The chamber consists of two primary parts: a support structure that forms the media and cell container, and a base that incorporates the desired PDMS formulation (Low, Mid, High), serving as the cell interaction interface (Figure 1B).

**Figure 1:**
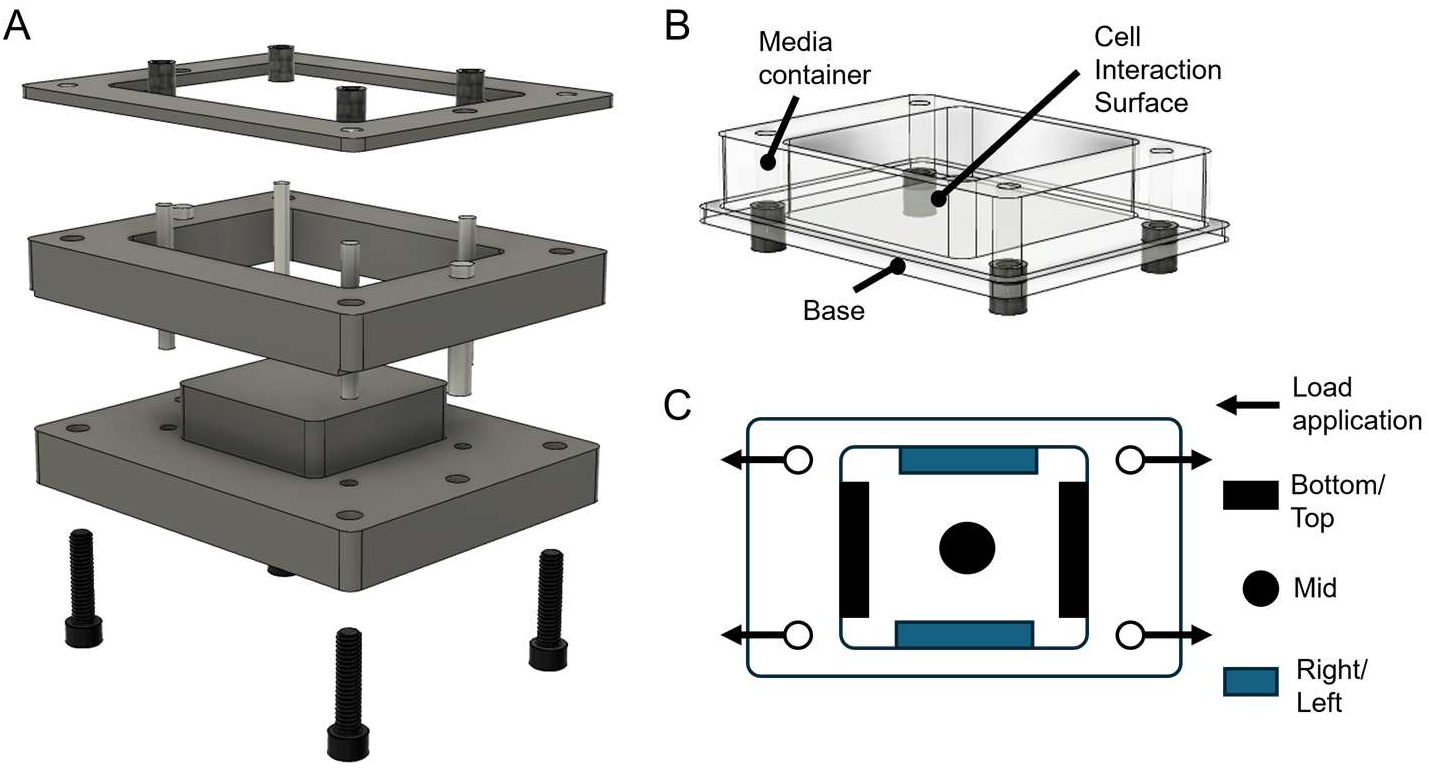
Design of the mold and custom stretching chamber. **A)** Exploded 3D model of the machined aluminum mold to fabricate cell stretching PDMS chambers. **B)** Model of the custom designed stretching chamber, composed of two parts: the support structure functioning as a media container, and the base structure being interchangeable with different stiffnesses for the cell interaction surface. **C)** Schematic of the top view of the stretching chambers showing the direction of applied strain and labeled regions for positional analysis.

For fabrication, the aluminum mold was cleaned with isopropanol (IPA, Sigma-Aldrich, 190764) and coated with Ease Release 200 (Mann) to minimize PDMS adhesion, according to the manufacturer’s instructions. To create the support structure (media container), a 50% wt blend of Sylgard 184 and Sylgard 527 was carefully poured into the mold up to the level of the cell interaction interface and pre-cured for 30 min at 65°C, allowing for effective bonding with the base PDMS layer. The mold was removed from the oven and the remainder (base) was filled up with the above-described Sylgard 184:527 ratio (Low, Mid or High) and further cured for 24 h at 65°C.

### 2.3 Tensile testing

Specimens for tensile testing of varying stiffness were molded into acrylic dog bone shaped samples according to ASTM D882-2010 standard (*n* = 5) and prepared from multiple production batches. For the commercial chamber dog bone shaped specimens (*n* = 4) were manually cut from the cell seeding area of the 10cc chamber (STREX).

The Young’s modulus of samples was determined via uniaxial tensile testing using a UniVert tensile tester equipped with a 1-10 N load cell (CellScale Biomaterials Testing). Each sample was preloaded with 0.1 N, and the samples were tested until rupture at a constant rate of deformation (10 mm/s). The Young’s modulus was calculated from the slope of the linear elastic region of the stress-strain curve.

### 2.4 Dynamic mechanical analysis (DMA)

DMA was completed on PDMS substrates using a TA Instruments HR20 rheometer with 8 mm cross-hatched upper and lower geometries. 8mm diameter samples (1 mm thickness) were stamped and oscillatory amplitude experiments (1 Hz, 20°C) were completed over the 0.1 to 40% strain range. Strain amplitudes ranging from 0.1 to 10% were shown to be in the linear viscoelastic region for all formulations. Accordingly, oscillatory frequency sweeps were run in triplicate from 0.1 – 30 Hz at 5% strain for 14 and 24% wt Sylgard 184/527 blends (Mid and High, respectively), as well as for pure Sylgard 184 (Commercial). Owing to the soft elastomeric nature of the Sylgard 527 (Low) substrate, oscillatory frequency sweeps were obtained from 0.1 – 30 Hz at 1% strain, also in triplicate. Additionally, oscillatory frequency sweeps were obtained at 37°C for all formulations.

### 2.5 Fourier Transform Infrared (FTIR) spectroscopy

A PerkinElmer Frontier FTIR spectrometer with an Attenuated Total Reflectance (ATR) attachment (diamond/ZnSe crystal) was utilized to analyze one sample per PDMS formulation (Low, Mid, High, Commercial). Spectra were measured in the 650–4000 cm⁻¹ range, averaging 16 scans per experiment.

### 2.6 Optical transparency

Optical transparency was measured using a UV-Vis spectrometer (Genesys 20, Thermo Scientific). The transmittance spectra of three PDMS samples were recorded over a wavelength range of 400 nm — 800 nm. One-millimeter-thick samples of the three different blends (Low, Mid, High) were evaluated and compared against standard micro cover glass slides (VWR) and commercially available microscope mountable chambers (SC-0022, STREX).

### 2.7 Chamber characterization using Digital Image Correlation (DIC)

Digital image correlation (DIC) was utilized to evaluate both axial and transverse average strains, and strain distributions on the cell seeding surface of the stretching chambers during uniaxial stretching. To enable strain measurement, a blacklight-sensitive ink (Millennium Colorworks, Ink Glow UV) was uniformly sprayed onto the surface to create a speckle pattern suitable for DIC imaging. Initially, the surface strain of the custom chambers was measured and compared to those of commercially available chambers across strain amplitudes ranging from 8% to 20% (*n* = 3). Subsequently, a detailed characterization was performed at 20% strain for the custom chambers (*n* = 7-8) and compared to the commercial chambers (*n* = 3, 10cc Chamber, STREX). Measurements were performed under sinusoidal uniaxial loading using the automated cell stretching system (STB-140-10, STREX).

First, a standard calibration procedure was performed using the DaVis Software (LaVision, 10.2.1). Subsequently, the chambers were imaged under cyclic sinusoidal uniaxial loading conditions at 1 Hz at varying strain amplitude using the StrainMaster Portable System (LaVision), including two charge-coupled (CCD) cameras (*f* = 50 mm lens) and two blue LED illuminations. Image sampling was done at 100 Hz. Data acquisition, image processing, and analysis were done within the software.

### 2.8 Bovine AF cell isolation and culture

Bovine tails were obtained from a local slaughterhouse. The tails were initially submersed in a betadine solution for disinfection and dissected under sterile conditions to separate the AF from the remaining zones of the IVD (NP, the transition zone NP/AF, outer AF border) using a surgical scalpel. The dissected AF was minced and subjected to a two-step enzymatic digestion process for cell isolation. Briefly, the AF tissue was incubated with 2.5 mg/mL of pronase (Sigma-Aldrich, 10165921001) in Dulbecco’s Phosphate-Buffered Saline (DPBS, Cytiva, SH30028.02) for 1 h at 37°C with 5% CO_2_ on a shaker following a 1 mg/mL of collagenase NB4 (Nordmark, S1745402) in HyClone Dulbecco’s Modified Eagle Medium (DMEM/F12, Cytiva, SH30023.FS) supplemented with 5% Fetal Bovine Serum (FBS, Cytiva, SH30396.03) and 1% Antibiotic Antimycotic solution (Anti-Anti, Cytiva, SV30079.01) incubation (37°C, 5% CO_2_) overnight on a shaker. The digested tissue was strained through a 70 µm sterile cell strainer (CELLTREAT, 229484), and the cell solution was then centrifuged for 10 min at 500 g. The enzyme solution was aspirated, and the cells were expanded in flasks coated with 10 µg/mL fibronectin (FN, EMD Millipore, FC010), using standard medium composed of DMEM/F12, 10% FBS, 1% Anti-Anti and 100 µM L-Ascorbic acid (AA, HelloBio, HB1238). AF cells were expanded at 37°C and 5% CO_2_, with medium changes three times per week, and passaged at 80% confluency.

### 2.9 PDMS surface modification and cell culture

PDMS-coated well plates and PDMS chambers were cleaned twice by immersing them in IPA and ultrasonicating for 5 min. Subsequently, they underwent an additional ultrasonic cleaning step in sterile DPBS to remove the excess IPA. PDMS substrates or chambers were then dried under UV light (30 min) in a sterile environment. Following drying, the samples were plasma-treated with a plasma cleaner (Harrick Plasma, PDC-001) before coating overnight in an incubator with 10 µg/mL FN for static cell experiments or with 50 µg/mL FN for stretching experiments to promote cell adhesion. AF cells (passage 2 to 4) were seeded into PDMS-coated plates, uncoated control plates (TCP), or cell stretching chambers. Cellular experiments were conducted at 37°C and 5% CO_2_.

### 2.10 AF cell viability and cytotoxicity assessment

AF cell viability (PrestoBlue^TM^ HS Cell Viability Reagent, Invitrogen, P50201) and cytotoxicity (CyQUANT™ LDH Cytotoxicity Assay Kit, Thermo Fisher Scientific, C20301) were used to determine cell attachment and proliferation over a 72 h period on PDMS substrates to indicate biocompatibility. Cell viability was also used to evaluate cellular responses to cyclic mechanical stretching.

Static conditions: To assess cell viability in PDMS, AF cells were seeded in 96-well plates covered with plasma-treated and FN-coated PDMS (3,000 cells/well) and incubated for 24 – 72 h at 37°C, 5% CO_2_. In a subset of wells, a collagen I coating (25 µg/mL, COL, Rat Collagen Type I, Sigma-Aldrich, C3867) was applied instead of FN for ECM coating comparison. Thereafter, the medium was replaced with a 1:10 dilution (1X, in DMEM/F12) of PrestoBlue™ and incubated for 3.5 h. A 24 h, an additional gentle wash with DPBS was performed prior PrestoBlue™ incubation to assess cell attachment on the PDMS substrate. Cells cultured on a regular 96-well plate (TCP) served as positive control and cells were lysed for negative control. In addition, cytotoxicity was tested in static conditions in which AF cells were cultured on PDMS coated surfaces in a 96-well plate, as above described, for 48 h and 72 h. Condition media was collected and LDH levels were measured according to the manufacturer’s instructions including and LDH lysis (positive) and untreated (negative) control.

Stretching conditions: To assess cell viability with stretching, AF cells were cultured on Mid stiffness stretching chambers (50,000 cells/chamber). After 48 h, the culture medium was replaced with serum-reduced medium (DMEM/F12, 5% FBS, 0.1% Anti-Anti, 100 µM AA). After 1 h of acclimatization, cells underwent cyclic sinusoidal stretching at 8% strain at 1 Hz for durations of 6 to 18 h, and medium was thereafter replaced with a 1:10 dilution of PrestoBlue™ reagent. Measurements were conducted as described above. Controls were maintained under identical conditions without mechanical stretching.

### 2.11 AF cell area and alignment under static and cyclic stretch conditions

For the cell area measurement, AF cells from each donor were individually cultured on plasma treated and FN coated PDMS formulations with varying stiffness levels (Low, Mid, and High) or untreated TCP as a control, in 6-well plates (5,000 cells/well). The medium was replaced after 48 h to serum-reduced medium containing 5% FBS, 0.1% Anti-Anti, and 100 µM AA, and cultured for an additional 24 h. For alignment studies under cyclic stretching, AF cells were cultured on Low or High stiffness chambers (50,000 cells/chamber). After 48 h, the culture medium was replaced with serum reduced media. Following a 1 h incubation, cells underwent cyclic sinusoidal stretching at 8% strain and a frequency of 1 Hz for 6 h.

For both cell area and alignment, cells were washed twice with DPBS and then fixed with 4% paraformaldehyde solution (PFA, Thermo Scientific, J19943-K2) for 15 min. Following fixation, the cells were permeabilized for 10 min with 1X PBS-Triton X-100 solution (Alfa Aesar, J63521). Thereafter, cells were blocked with 4% bovine serum albumin (BSA, VWR, AAJ64100-09) in DPBS for 30 min and incubated with Alexa Fluor™ 594 phalloidin (1:400 dilution, Invitrogen, A12381) in 4% BSA for another 30 min. The nuclei were stained with Hoechst dye (1:3000 dilution, Thermo Scientific, H3570) in DPBS for 5 min. Finally, the cells were washed and covered with DPBS for imaging.

For each condition, three locations were imaged with an Olympus IX81 microscope equipped with an ORCA-Flash 4.0LT + camera. Images were processed using FIJI (ImageJ) software, and the cell area and alignment were quantified using CellProfiler (4.2.8). For alignment visualization, plots were created using Matlab (R2023a).

### 2.12 Statistical analysis

Data were checked for consistency, and normality was assessed using a Shapiro-Wilk test. Bootstrap-ANOVA was used to test effects globally and pairwise testing was done using unpaired t-tests with Welch’s correction and bootstrap-t tests. Bootstrap-t tests were used due to the small sample size and potential violation of normality. The method generates a simulated sampling distribution from the sample itself, offering a more reliable p-value and confidence interval when parametric assumptions are questionable. All tests were two-sided, and p-values < 0.05 were considered statistically significant. To statistically assess the cellular alignment, differences in their distribution profile were tested using a Kolmogorov-Smirnov test. Moreover, a linear regression between engineering and measured strain was applied to obtain R^2^ values. Data are presented as mean ± standard deviation (SD). Normality and t tests were conducted using GraphPad Prism (10.2.0), bootstrap t tests and Kolmogorov-Smirnov tests were done in Mathematica 13 (Wolfram Research, Inc. Champaign, Illinois, 2021).

## 3 Results

### 3.1 Mechanical characterization of the PDMS substrates revealed a broad range of mechanical properties

The PDMS substrates were fabricated by adding Sylgard 184 and Sylgard 527 at mass ratios of 0%, 14%, 24% or 100% Sylgard 184 relative to Sylgard 527 (as detailed in Table 1) and cured at 65°C for 24h. These formulations are herein referred to as Low, Mid, and High (custom) and commercial respectively.

Tensile testing demonstrated a clear increase in Young’s modulus with higher Sylgard 184 content (Figure 2A). Statistical analysis confirmed significant differences among the formulations (*p <* 0.001), with moduli ranging from 8.72 ± 3.09 kPa for the Low formulation to 238.00 ± 31.20 kPa for the High formulation (Table 2). Since these formulations were eventually incorporated into custom stretching chambers, Young’s modulus of the commercial STREX stretching chamber was measured to set a benchmark. The test revealed a modulus of 1217.00 ± 137.50 (Figure 2A), which is substantially higher than the highest substrate stiffness (*p <* 0.001).

**Figure 2:**
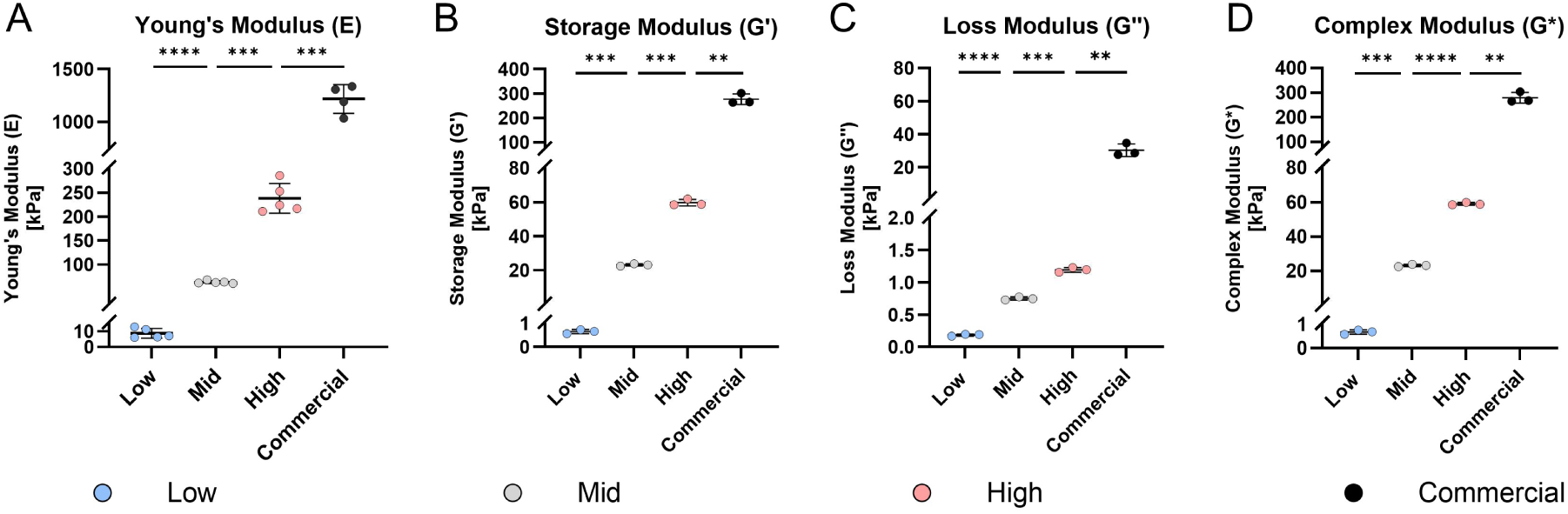
Mechanical characterization of the PDMS formulations prepared with varying ratios of Sylgard 184 to Sylgard 527 (0, 14, 24 or 100% Sylgard 184). **A)** Stiffness (Young’s modulus) of the PDMS substrates (Low, Mid, High, *n* = 5), determined by tensile testing, and compared to PDMS substrate of commercial chambers (commercial, *n* = 4). **B-D)** Rheological properties of PDMS formulations (Low, Mid, High, *n* = 3), and compared to commercial formulation (Sylgard 184, *n* = 3), assessed by dynamic mechanical analysis (DMA) at 1 Hz, including storage modulus **(B)**, loss modulus **(C)**, and complex modulus **(D)**. Data are presented as mean ± SD. ** *p* < 0.01, *** *p* < 0.001, **** *p* < 0.0001.

**Table 2:**
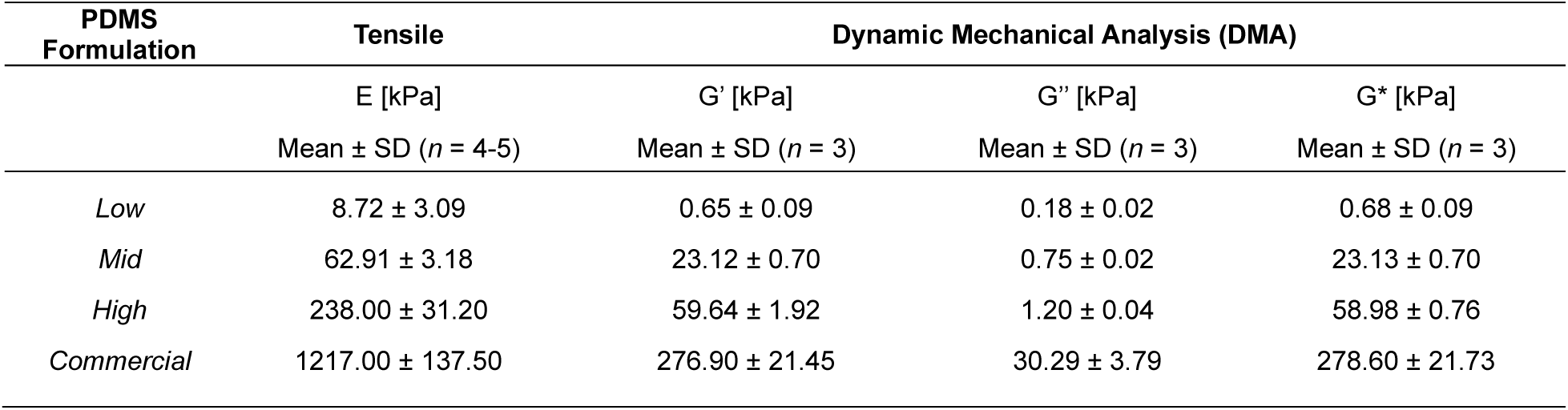
Tensile and rheological properties of the PDMS formulations, including Young’s- (E), storage- (G’), loss- (G’’), and complex (G*) modulus at a selected frequency of 1 Hz. Values represent mean ± SD.

In addition to tensile testing, DMA was conducted to characterize the viscoelastic properties of the PDMS formulations (Figures 2B-D). To determine the linear viscoelastic range, an amplitude and frequency sweep were performed. The amplitude sweep (Supplementary Figure 1A-C) indicated linear viscoelastic behavior between 0.8 to 6.3 strain percentage, while the frequency sweep (Supplementary Figure 2) showed a stable response between 1 and 10 Hz. Based on these results, a physiological testing frequency of 1 Hz was selected to represent storage modulus (G’), loss modulus (G’’), and complex modulus (G*) (Figures 2B-D).

DMA confirmed that increases in the amount of Sylgard 184 resulted in commensurate increases in the shear storage, loss, and complex moduli (G’, G’’ and G* respectively) of substrates. As summarized in Table 2, significant differences (*p <* 0.01) were found for storage moduli G’ between groups, ranging from 0.65 ± 0.09 kPa (Low) to 59.64 ± 1.92 kPa (High), while reaching 276.90 ± 21.45 kPa for the commercial formulation (Sylgard 184 only) under the same curing conditions. Although the Loss modului G″ rose with increasing concentration of Sylgard 184, it remained relatively low across all blends (0.18 ± 0.02 to 1.20 ± 0.04 kPa), while reaching 30.29 ± 3.79 kPa in the commercial formulation, with statistical significance between all groups (*p* < 0.01). The Low formulation exhibited the highest relative viscous contribution (G″/G′ = 0.28). Furthermore, DMA performed at 37 °C indicated that the mechanical properties of each formulation were stable at physiological temperature (Supplementary Figure 1D-F).

### 3.2 Optical and chemical PDMS properties

High quality imaging requires highly transparent substrates. PDMS is widely recognized for its excellent transparent properties and usage is reported even in optical applications [36]. Figure 3A quantitatively compares the transmittance of our three PDMS formulations (Low, Mid, High) to microscope glass slide and commercial STREX chamber. All formulations demonstrated excellent optical properties in the visible spectrum (400-800 nm), with average transmittance exceeding 90% and a trend in higher transmittance observed for increasing Sylgard 184 components. The custom substrates matched or exceeded the transmittance of microscope glass slides and commercial chambers. Although minor differences were detected between formulations, these are unlikely to affect routine cell-imaging routine cell-imaging experiments.

**Figure 3:**
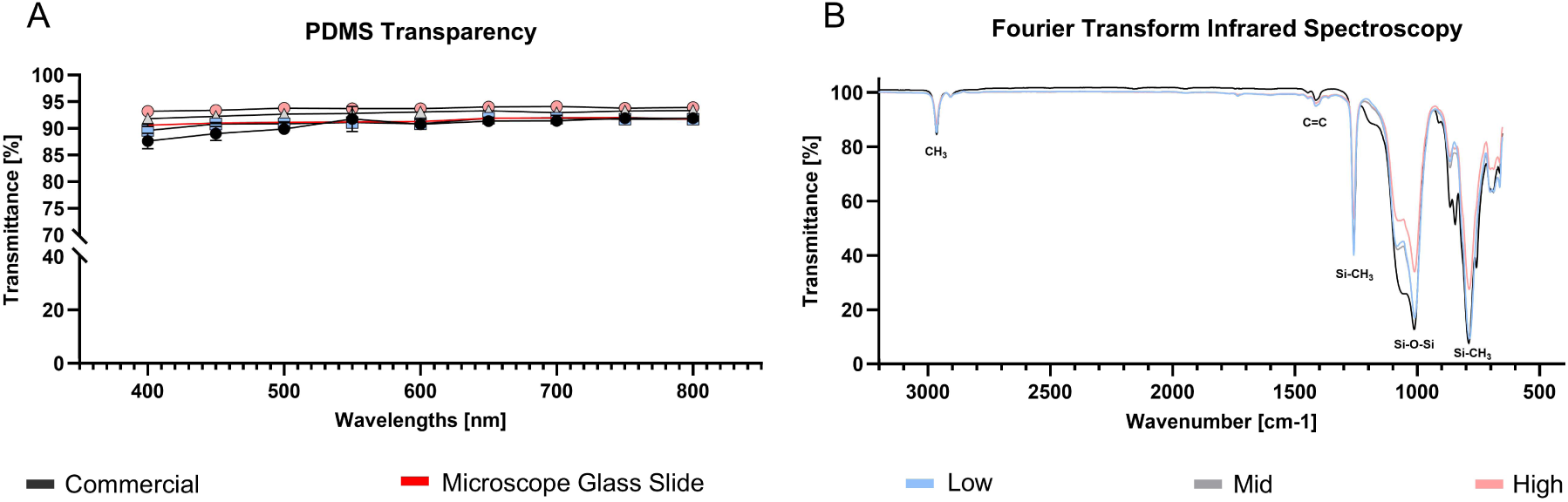
Optical and chemical PDMS properties. **A)** Optical transparency of PDMS substrate (Low, Mid, High) compared to commercial microscope mountable stretching chambers (STREX) and microscope glass slide. Data include mean ± SD, (*n* = 3). **B)** FTIR-ATR spectra, represented in transmittance, of cured PDMS substrates (Low, Mid, High, commercial, *n* = 1).

As shown in Figure 3B, all PDMS formulations exhibit characteristic infrared peaks as described by Johnson *et al.* [37]. This includes peaks at 786–791 cm^-1^ (related to −CH_3_ rocking and Si-C stretching in Si-CH_3_), 1010–1080 cm^-1^ (related to Si-O-Si stretching), 1257-1260 cm^-1^ (related to CH_3_ deformation in Si-CH_3_), and 2964-2965 cm^-1^ (related to asymmetric CH_3_ stretching in Si-CH_3_). As expected, all formulations show nearly similar profiles.

### 3.3 Biocompatibility assessment and static AF cell responses on PDMS formulations

PDMS substrates are known to be biocompatible [26], however, studies on formulations including Sylgard 527 are limited. Therefore, Figures 4A-E focuses on cellular viability and cytotoxicity of the fabricated PDMS formulations over 72 h (Figure 4A). All PDMS formulations supported AF cell attachment and growth with no significant differences in viability at 24 h and 48 h across all conditions and proliferation occurring between 48 h and 72 h. The cell viability at 72 h suggests that the High stiffness substrates (10,589.9 ± 1,004.6 fluorescent units) was significantly lower than on the uncoated TCP control (12,923.5 ± 522.0 fluorescence units, *p <* 0.05), and also lower than the Low substrate (13,178.4 ± 376.8 fluorescent units, *p* < 0.05). The viability of cells on the Low substrate was higher than on the Mid substrate (11,423.7 ± 512.6 fluorescent units, *p* < 0.05).

**Figure 4:**
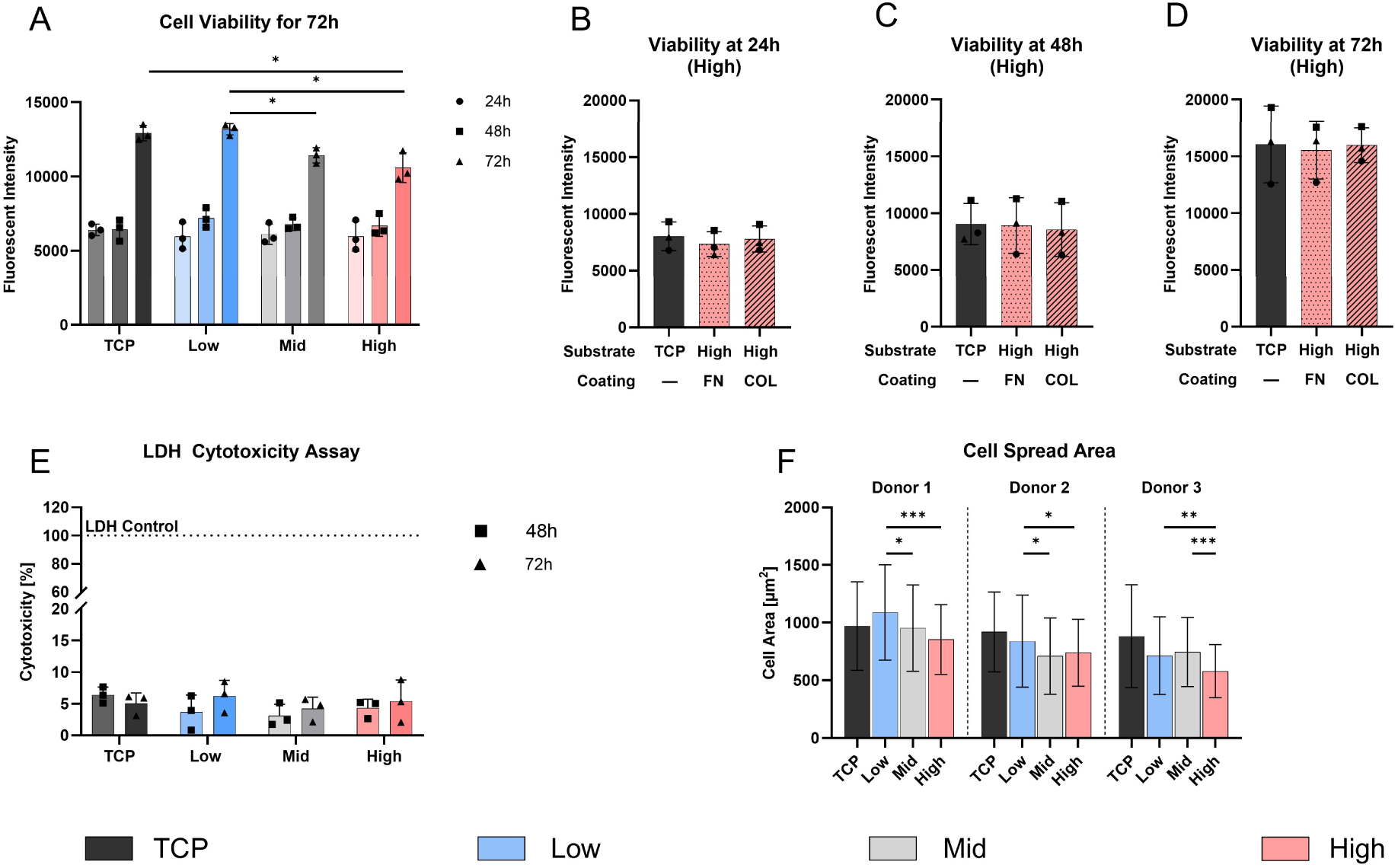
Biocompatibility evaluation and substrate interaction on PDMS formulations over 72 h. **A)** AF cell viability on PDMS substrates (Low, Mid, High) and TCP, measured via PrestoBlue^TM^ assay at 24, 48, and 72 h. **B-D)** Cellular viability on High PDMS substrates coated with Fibronectin (FN) and Collagen 1 (COL), compared to uncoated TCP as a control. **E)** Cytotoxicity (LDH assay) evaluation at 48 h and 72 h of all substrates, including TCP and LDH control. **F)** Quantification of cell spread area (µm^2^) on various PDMS formulations and TCP after 72 h, with images captured at three random locations per condition. Data represents mean ± SD from three individual donors. * *p* < 0.05, ** *p* < 0.01, *** *p* < 0.01.

To assess surface coating effects, the High PDMS substrate was functionalized with fibronectin (FN) and collagen I (COL). At all time points (24, 48, 72 h), viability on FN and COL coatings was comparable to uncoated TCP (15,990.5 ± 1526.8 fluorescent units), with no significant differences between groups (Figures 4B-D) indicating equal cell growth and attachment.

To assess any potential cytotoxic effects of the PDMS substrates, the LDH release of AF cells was measured at 48 h and 72 h. The results indicated minimal cytotoxicity with no significant differences among all conditions (Figure 4E).

Furthermore, the cell spreading area was determined after 72 h by fluorescent staining of actin and cell nuclei (Figure 4F). Donor 1 cells exhibited the largest mean cell spreading area on the Low stiffness substrates (1,090.0 ± 413.2 µm^2^), which decreased with increasing substrate stiffness, with significant differences to the Mid (953.4 ± 374.4, *p* < 0.01) and High (853.9 ± 303.3 µm^2^, *p* < 0.001) substrate. Although donor 2 and 3 showed similar trends, stiffness-dependent patterns were less consistent: Donor 2 showed a slight significance in Low (840.0 ± 399.1 µm^2^) compared to both, Mid (710.1 ± 330.7 µm^2^, *p* < 0.01) and High (739.7 ± 290.2 µm^2^, *p <* 0.01) substrates, while donor 3 showed significant differences between Low and High formulations (*p* < 0.01).

### 3.4 Strain characterization of stretching chambers under mechanical loading using digital image correlation

Custom PDMS stretching chambers with integrated stiffness were fabricated using the developed aluminum mold. Given the uniqueness of this approach, and the absence of direct reference data, we characterized the strain on the cell interaction surface to identify differences among the custom and commercial chambers and to investigate the strain transmitted by our stretching chambers to AF cells during stretching. The results were compared to the commercially available stretching chambers from STREX. To achieve this, the study used DIC, an optical and non-contact method, that measures strains over a large field of an elastic surface by tracking a randomly applied speckle pattern during deformation [38]. We applied a range (8 to 20%) of engineering strains (ε*_eng_*) to the chambers using the STREX bioreactor, at a physiological frequency of 1 Hz.

PDMS chambers exhibited a strong linear relationship between the applied ε*_eng_* and the measured average maximum axial strain on the cell interaction surface for both, commercial chambers (Figure 5A, R^2^ = 0.995) and across all custom stretching chambers levels (Figure 5E, R^2^ ≥ 0.974). The axial strain on the cell interaction surface was accompanied by a transverse strain component, perpendicular to the main loading axis (axial). Similarly, Figure 5B and 5F displayed that the absolute transverse strain increased linearly with the applied strain for all chambers (R^2^ = 0.993 for commercial chambers and R^2^ > 0.965 for custom chambers). Statistical analysis revealed significant differences in both axial (*p* < 0.001) and transverse strain (*p* < 0.05) measurements across all applied strain levels in commercial chambers (Figures 5C and 5D) as well as across all strain levels within a stiffness level in axial (*p* < 0.05) and transverse for custom chambers (Figures 5G and 5H). Moreover, Figures 5G and 5H display significant differences in majority of measurements across the stiffness levels, with only a few exceptions.

**Figure 5:**
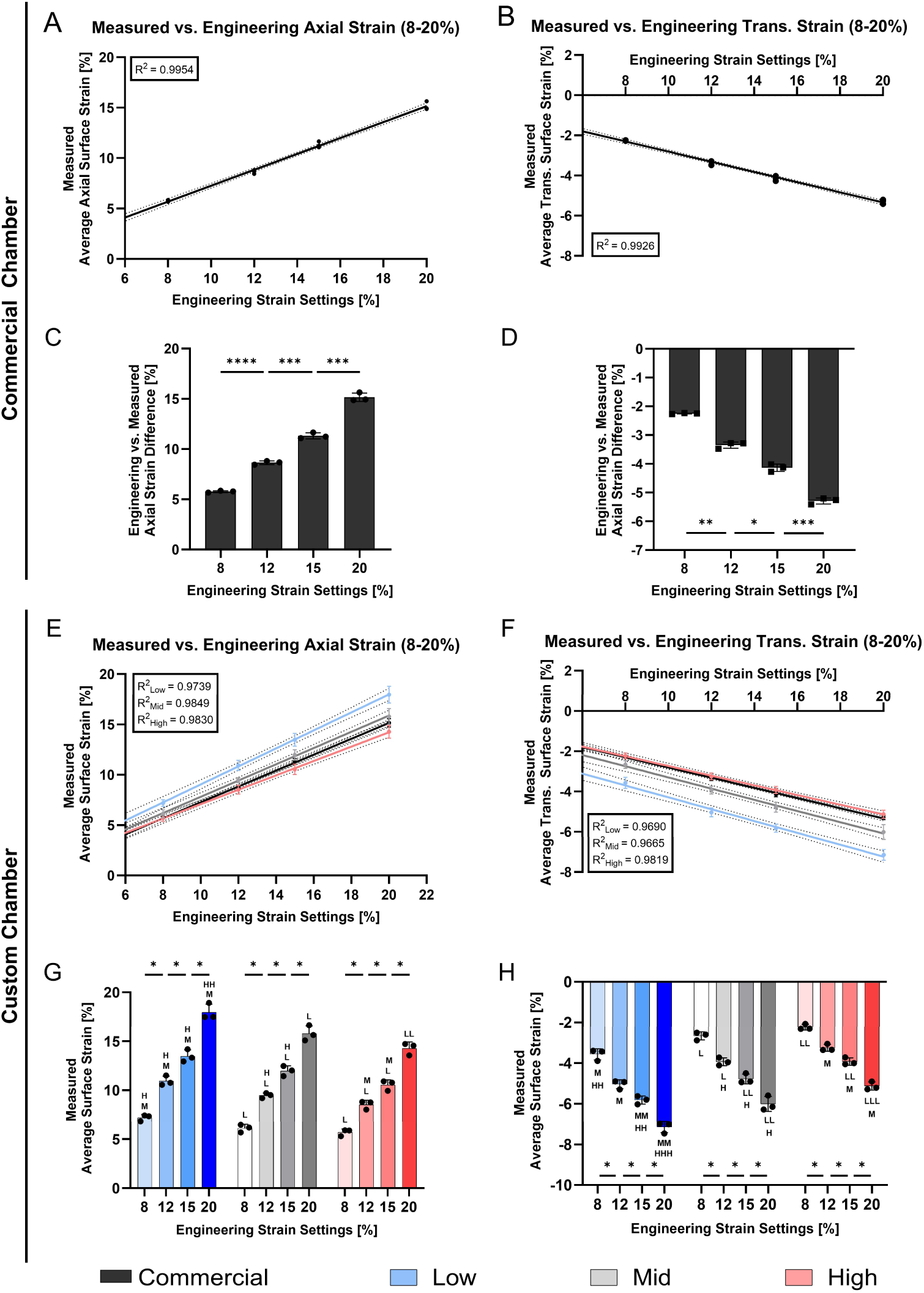
Average surface strain measurement on the cell interaction surface of commercial (STREX) and custom stretching chambers. Strain measurements were performed across engineering strain settings ranging from 8% to 20%. Engineering strain was applied using the commercially available STB-140-10 system from STREX. **A,B,E,F)** Linear regression analyses of average axial **(A,E)** and transverse strain measurements **(B,F)** of commercial **(A,B)** and custom stretching chambers **(E,F)**. **C,D,G,H)** Bar plot of the average axial **(C,G)** and transverse strain measurements **(D,H)** in commercial chambers **(C,D)** and custom stretching chambers **(G,H)**. Linear regression represents three individual replicates with indicated error lines. Bar plots indicate mean ± SD, (*n* = 3). Statistical significance is indicated as follows: ^L, M, H^ indicates significance among stiffness levels (Low, Mid and High respectively) within equal strain levels where * indicates significance among strain levels within a strain level, * *p* < 0.05, * *p* < 0.05, ** *p* < 0.001 (* also representative for ^L, M,H^).

Finally, the difference of the applied ε*_eng_* and the measured average maximum axial strain were compared (Supplementary Figure 3). In commercial chambers, a difference of 2.24 ± 0.14% was found at an applied ε*_eng_* of 8%. The difference increased to 5.02 ± 0.23% at 20% ε*_eng_*. This trend was also observed in custom chambers. Low stiffness chambers showed smallest differences (0.80 ± 0.33% at 8% ε*_eng_*, 2.04 ± 0.83% at 20% ε*_eng_*), while High stiffness chambers showed the largest discrepancies (2.30 ± 0.34% at 8% ε*_eng_*, 5.72 ± 0.64% at 20% ε*_eng_*).

We further quantified average axial and transverse strains at the loading peak on the cell interaction surface under a 20% ε*_eng_* (Figures 6A,B). Figure 6A illustrates significant differences in axial strain (*p* < 0.01) across all chamber types apart from the commercial and Mid chamber and significance in transverse strain measurements (*p* < 0.01) at the cell interaction surface across all chamber types. At an applied engineering strain of 20%, the commercial chamber exhibited an average axial strain of 15.15 ± 0.42% and a transverse strain of −5.29 ± 0.11%. Among the custom chambers, the Low stiffness showed the highest axial strain (18.16 ± 0.57%) and the most negative transverse strain (−7.20 ± 0.33%). This was followed by the Mid stiffness chamber with an axial strain of 15.25 ± 0.89% and a transverse strain of −5.98 ± 0.38%, and the High stiffness chamber with the lowest axial strain (13.06 ± 0.90%) and lowest negative transverse strain (−4.85 ± 0.31%).

**Figure 6:**
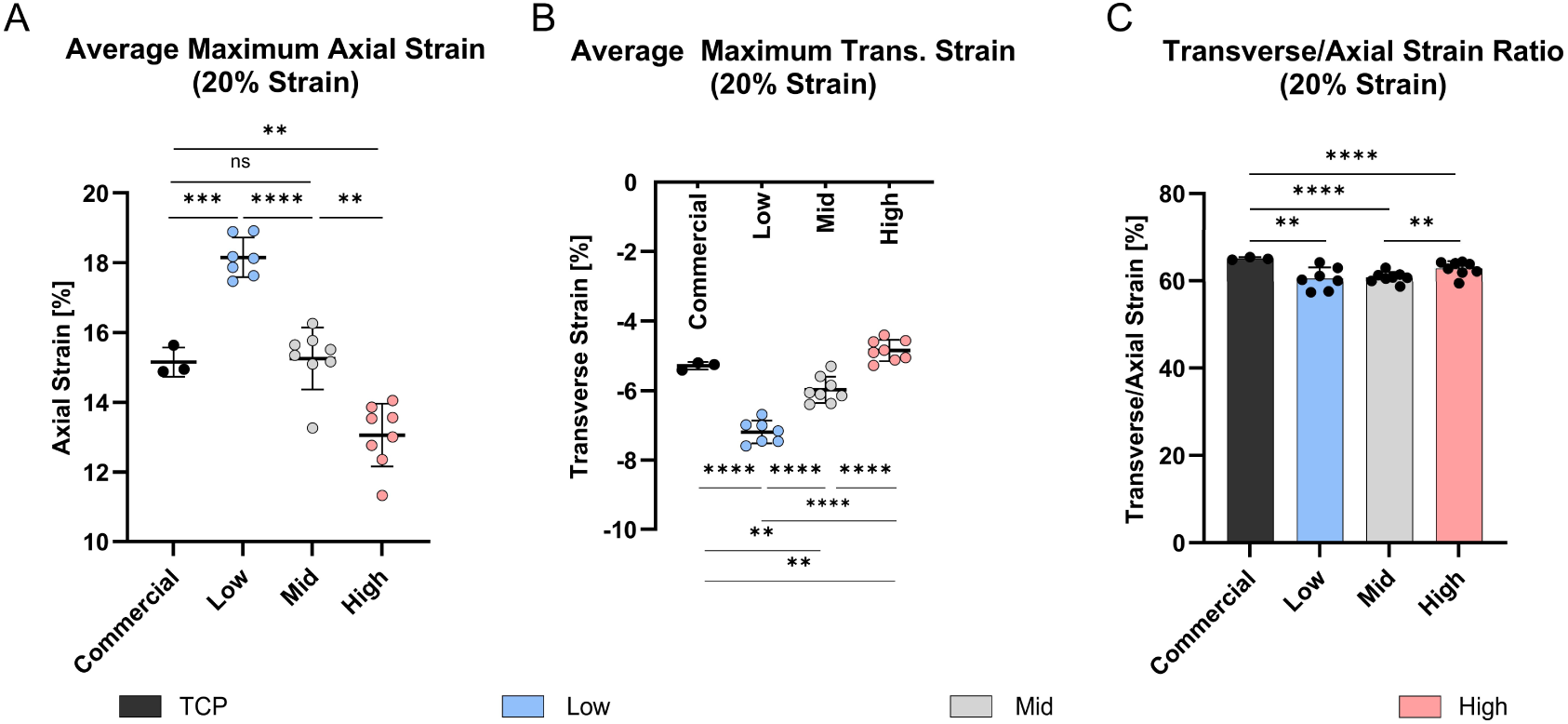
Average maximum strain at the cell interaction surface of PDMS chambers at 20% and 1 Hz deformation. **A)** Average axial and **B)** transverse maximum strain for commercial and custom PDMS chambers. **C)** Relative major axis (axial) to minor axis (transverse) strain in percentage (((ε_trans_/ε_axial_) +1) x100). (commercial: *n* = 3, custom: *n* = 7-8), mean ± SD, *ns* = non-significant, * *p* < 0.05, ** *p* < 0.01, *** *p* < 0.001, **** *p* < 0.0001.

Stretching the PDMS chambers in a specific direction is expected to lead to predominant strains along the main loading axis. Due to the acting transverse strains, we aimed to quantify the relative average contribution of the major loading axis (axial) to the transverse axis strains in percentage. In the commercial chamber, 65.09 ± 0.29% of the total strain was aligned with the loading direction, while the remainder was attributed to transverse deformation. Significant differences were observed when comparing the commercial chamber to the custom chambers (*p* < 0.01), which exhibited axial strain contributions of 60.51 ± 2.55% (Low), 60.79 ± 1.22% (Mid) and respectively 62.82 ± 1.68% (High) (Figure 6C).

In addition, the mechanical behavior of the chambers during the entire deformation cycle (1 Hz, 20%) was quantified on representative samples. The resulting average axial strain curves, over one deformation cycle, showed a clear sinusoidal pattern for both the custom and commercial chambers. No obvious differences were observed across stiffnesses, with a strain peak around 500 ms and a full cycle completion at 1 s (Figures 7A,B). Notably, the cyclic deformation pattern of the commercial chamber closely resembles that of the Mid stiffness chamber. Figures 7C and 7D presents representative axial strain distribution images over the entire deformation cycle. The commercial chamber (Figure 7C) and the custom chambers (Figure 7D) displayed a characteristic axial strain pattern on the cell interaction surface, most prominent at the peak of the loading. In all designs, the highest axial strain values were localized in proximity to the load application points, and extending to the middle, while the lowest strain values were observed at the lateral side of the membrane (right and left side of the chamber, Figure 1C). High and Mid stiffness exhibited comparatively more uniform strain distributions across the surface than Low stiffness chambers (Figure 7D), reflected in lower mean SDs in Table 3, or represented as a line graph across the x-Axis in Supplementary Figure 4.

**Figure 7:**
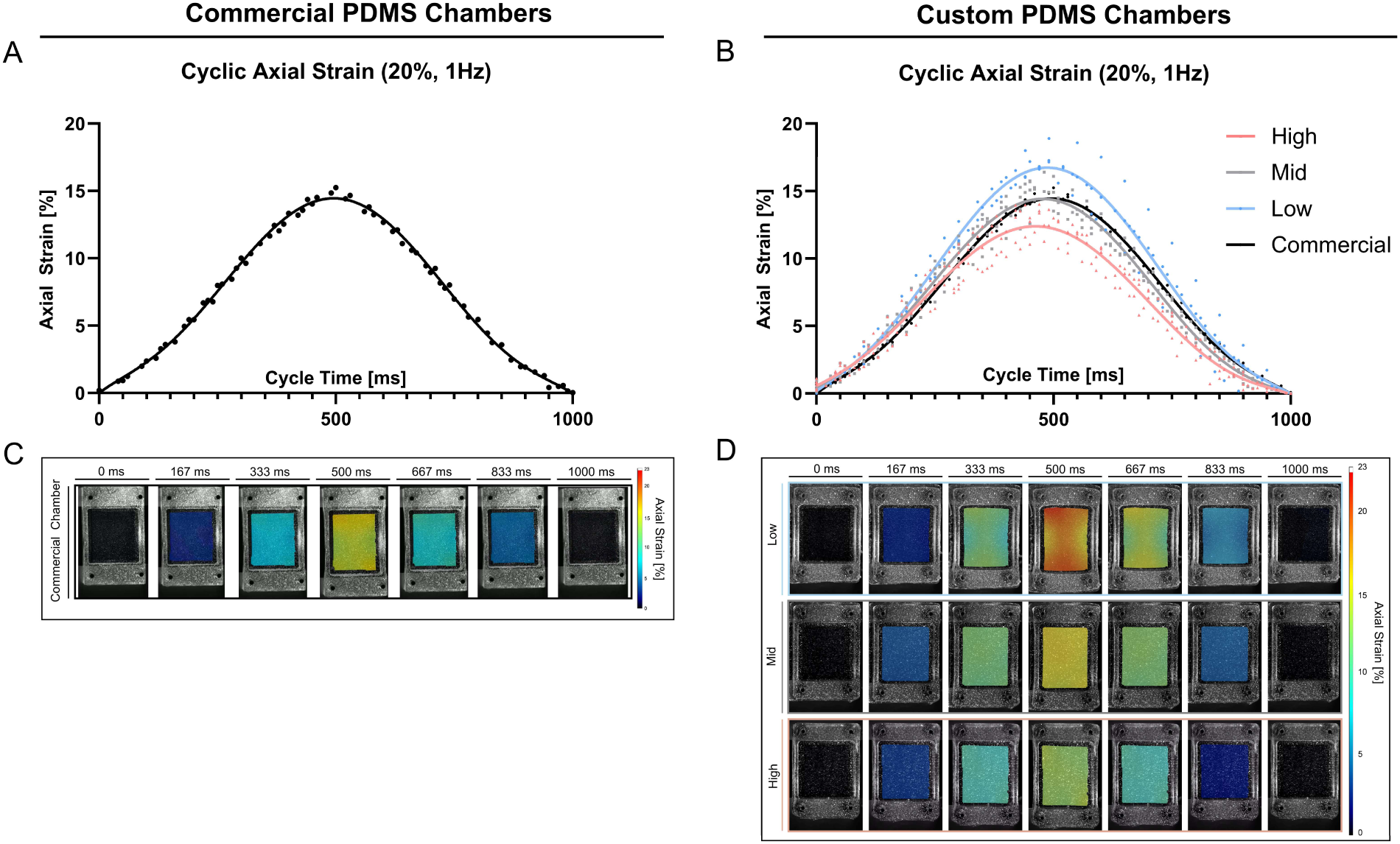
Average cyclic axial strain measurements on PDMS cell surface interaction at 1 Hz and 20% strain. **A,B)** Measured average axial strain within one deformation cycle of 1 Hz (0-1000 ms) in commercial stretching chambers **(A)** and custom stretching chambers **(B)**. Data represents three individual replicates, and a six-order polynomial curve was fit, R^2^ > 0.95. **C,D)** Axial strain distribution at the cell interaction surface (*representative images*) within one deformation cycle for commercial chambers **(C)** and custom chambers **(D)**.

**Table 3:** Axial and transverse strain distributions were calculated by averaging the strain distribution measurements from each individual chamber from the entire dataset. Variability across chambers is represented by the mean standard deviation (SD) and range.

|  | Axial |  |  |  |
| --- | --- | --- | --- | --- |
|  | Commercial | Low | Mid | High |
| Mean Range [%] | 3.02 | 9.33 | 4.82 | 5.23 |
| Mean SD [%] | 0.65 | 1.79 | 0.82 | 0.89 |
|  | Transverse |  |  |  |
|  | Commercial | Low | Mid | High |
| Mean Range [%] | 3.05 | 3.63 | 4.55 | 5.67 |
| Mean SD [%] | 0.73 | 0.81 | 0.92 | 1.09 |

Following the axial strain analysis, the cyclic transverse deformation of the PDMS chambers was quantified. As shown in Figures 8A and 8B, the deformation curves also exhibited a sinusoidal pattern, with maximum negative transverse strain at approximately 500 ms, consistent with the timing of axial strain. As expected, the full deformation cycle ranges 1 s at an applied frequency of 1 Hz.

**Figure 8:**
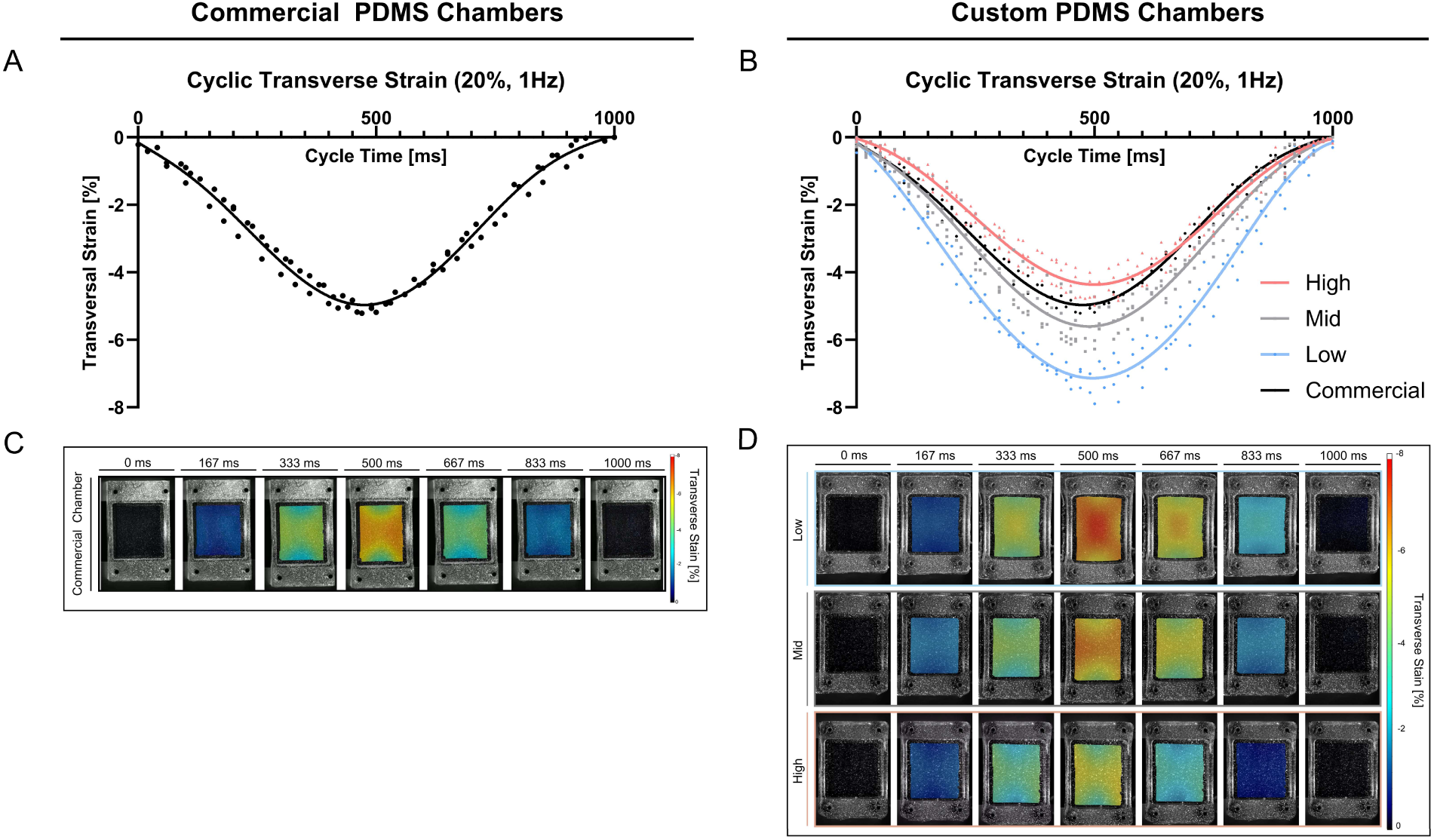
Average cyclic transverse strain measurements on PDMS cell surface interaction at 1 Hz and 20% strain. **A,B)** Measured average transversestrain within one deformation cycle of 1 Hz (0 – 1000 ms) in **A)** commercial stretching chambers and **B)** custom stretching chambers. Data represents three individual replicates, and a six-order polynomial curve was fit, R^2^ > 0.95. **C,D)** Transverse strain distribution (*representative images*) at the cell interaction surface within one deformation cycle for **C)** commercial chambers and **D)** custom chambers.

Figures 8C and 8D display representative transverse strain distribution maps over the entire deformation cycle. These maps reveal that the highest absolute transverse strain values are observed at the lateral edges (left and right, Figure 1C) of the chambers, extending inward towards the center. In contrast to the axial strain behavior, the lowest absolute transverse strain magnitudes occurred between the regions of load application (bottom and top), where the axial strain peak was previously seen. Moreover, representative images of the Poisson’s ratio distribution demonstrate the relative contribution of the transverse strain to the axial strain at the cell surface revealed highest relative contributions at the lateral side of the chambers extending toward the center, and lowest contributions at the top and bottom of the chamber between the load application points (Supplementary Figure 5).

Figure 9 quantitatively represents axial and transverse strain distributions at 20% applied engineering strain in three representative samples per condition, whereby measurements at a pixel-level are displayed as histograms. The entire dataset is shown as a boxplot in Supplementary Figure 6 or as individual values (mean, SDs) in Supplementary Table 1. Strain variability, expressed as the mean and standard deviation of individual chamber strain distributions, is summarized in Table 3.

**Figure 9:**
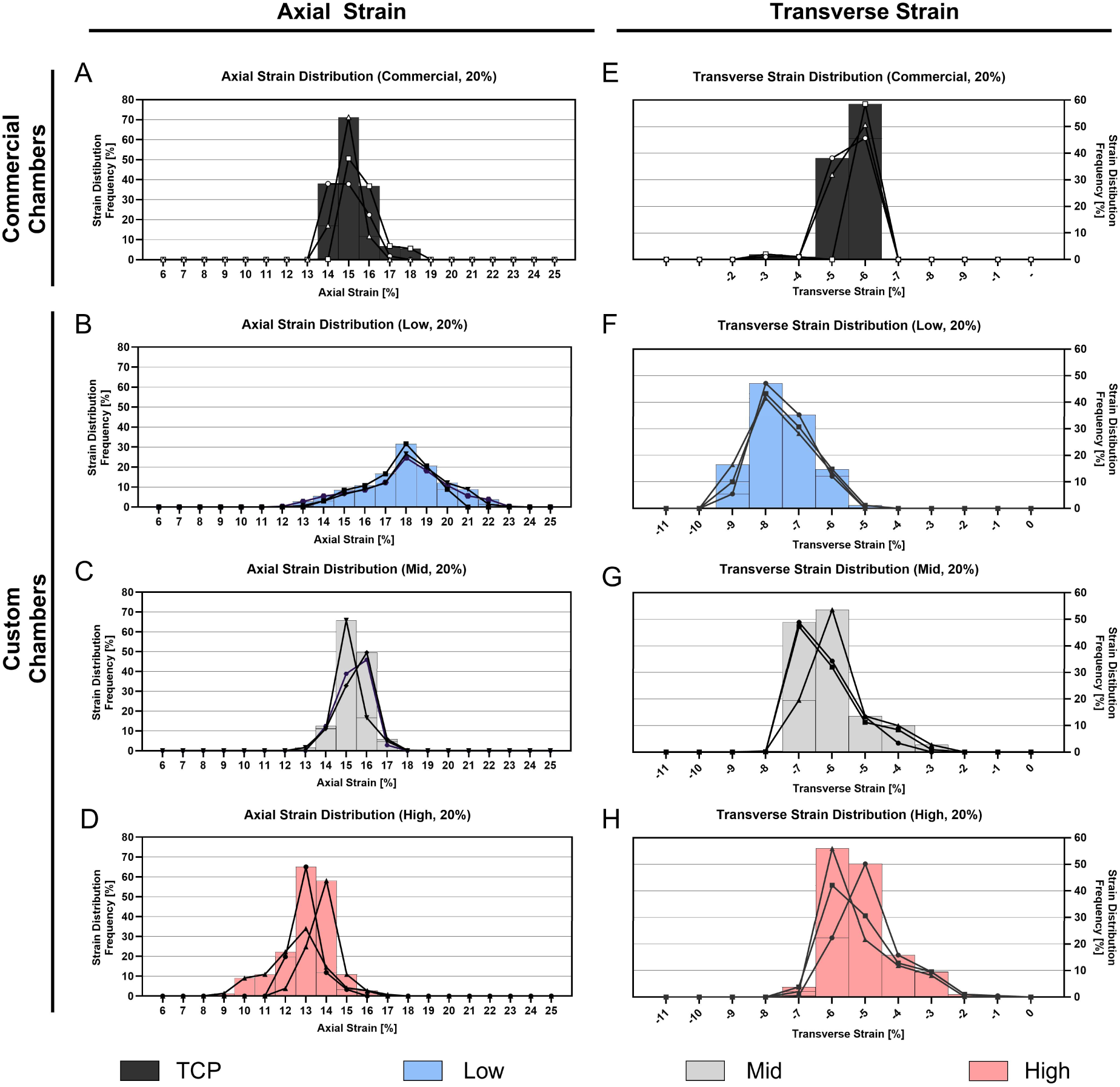
Frequency distribution plot of individual strain distribution characterization within a stretching chamber at maximum engineering strain settings of 20%. Data includes three representative samples. **A-D)** Axial strain distribution within the cell interaction surface of the PDMS chambers for commercial **(A)** and custom chambers **(B-D)**. **E-H)** Transverse strain distribution within the cell interaction surface of the PDMS chambers for commercial **(E)** and custom chambers **(F-H)**.

**Figure 10:**
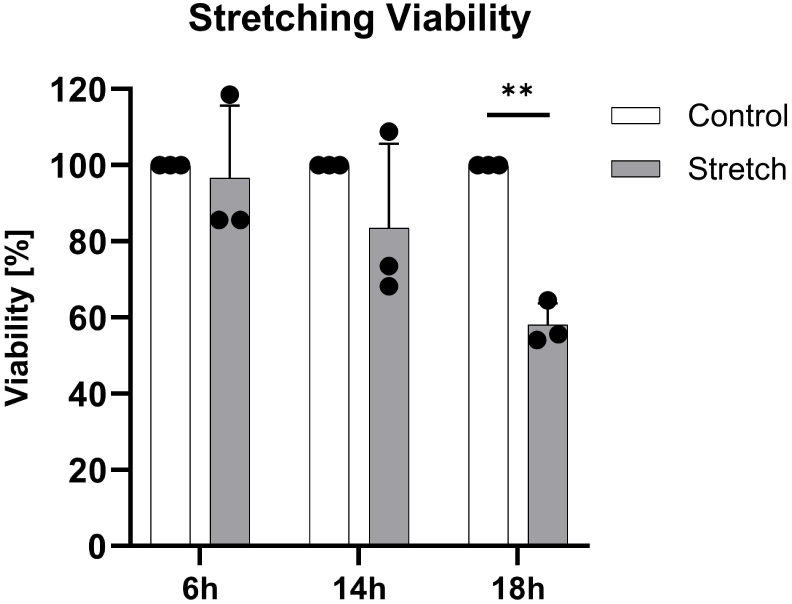
Annulus fibrosus (AF) cell viability over 6 h, 14 h and 18 h of dynamic stretching on Mid stretching chambers (8%, 1 Hz). Data represents mean ± SD, (*n* = 3). ** *p* < 0.01.

At an applied engineering strain of 20%, the commercial chamber (Figure 9A) exhibited the highest percentages of axial strains at 15%, with the lowest mean variability (3.02 ± 0.65%, Table 3) compared to custom chambers. In contrast, Low-stiffness chambers reached lower percentages of strain pixels at the peak of 18% but were accompanied by greater variability (9.33 ± 1.79%) in strain distribution. As chamber stiffness increased, peak axial strain decreased. Mid and High-stiffness chambers showed comparable axial strain distributions (Figures 9C,D), as reflected in their similar standard deviations (Table 3). A similar pattern was observed for transverse strain, where the commercial chamber had the narrowest range (3.04 ± 0.73%), while the High-stiffness chamber exhibited the widest range (5.23 ± 0.89%). These results suggest that increased chamber stiffness leads to reduced peak strains but increased variability in strain distribution.

### 3.5 Dynamic AF cell responses on PDMS stretching chambers

Cyclic mechanical loading creates a challenging environment for cells, potentially compromising their viability or attachment over time. Figure 9 assessed cell viability in Mid stiffness chambers subjected to dynamic stretching. After 6 h of loading, viability remained high at 96.62 ± 19.02%. However, prolonged exposure led to a decline of 83.56 ± 22.09% viability at 14 h and a significant reduction (*p* < 0.01) to 58.12 ± 5.64% at 18 h.

To assess the impact of load transmission and substrate stiffness on cell alignment, we analyzed cellular orientation (Figure 11), on Low and High stiffness chambers, representing the two extreme conditions of our chambers. Under static conditions, cells showed no distinct alignment, whereas after 6 h of cyclic loading at 1 Hz, generally a significant cell alignment, perpendicular to the stretch direction across all regions and stiffness was observed (*p* < 0.0001). Although, loaded cells on the Low substrate significantly aligned, the degree was lower than in the High stiffness chambers (*p* < 0.05). The degree of alignment also varied by location (imaging locations indicated in Figure 1C) with a significant difference in the alignment at: Top/Bottom and Mid locations (Figures 11B,C,F,G) compared to the alignment pattern at Right and Left areas (*p* <0.0001, Figure 11D,H).

**Figure 11:**
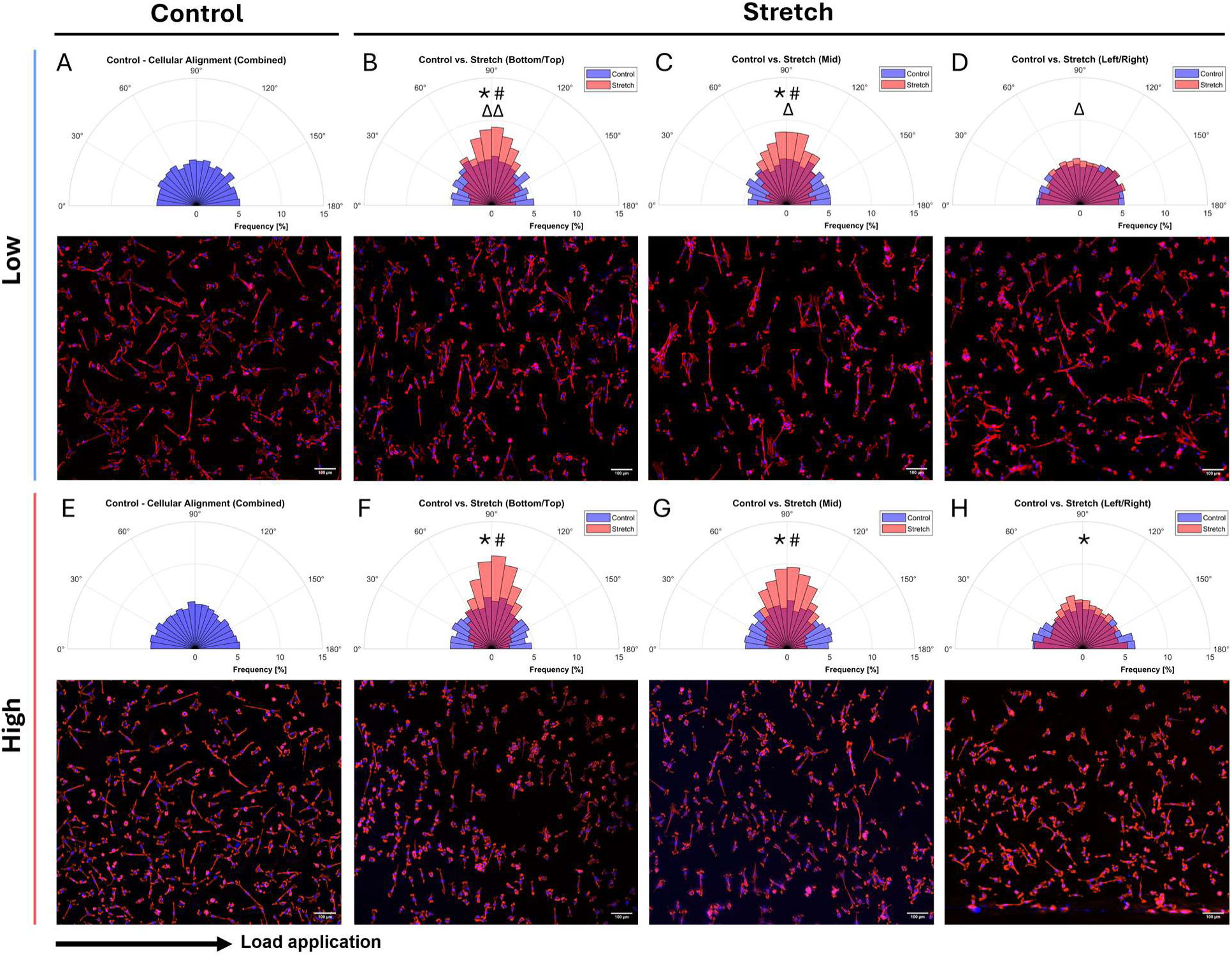
Cellular alignment (=90°) of static (**A,E**) and stretched (**B,C,D,F,G,H**) samples (8%, 1 Hz) in Low (**A-D**) and High (**E-H**) stiffness chambers, shown as polar histograms and representative fluorescent images. Images were taken from different strain regions within the chamber, including Top/Bottom **(B,F)**, Mid **(C,G)** and Left/Right **(D,H)**. Panel A and E display data from all combined locations in the static (control) condition with one representative image. For each location, data was obtained from three donors, with one representative image each. The arrow indicates the direction of load application. * Indicates significance: control vs. stretch (*p* < 0.0001). ^#^ Indicates significance: stretch Bottom/Top, Mid vs. stretch Right/Left (*p* < 0.0001) and ^Δ^ indicates significance: Low vs. High (^Δ^ *p <* 0.05, ^ΔΔ^ *p* < 0.01).

## 4 Discussion

Mechanical cues, particularly ECM stiffness and mechanical loading, play pivotal roles in IVD disease progression by compromising the structural integrity of the AF. While both factors have been individually studied in the context of IVD degeneration [14, 39, 40], their combined effect remains underexplored. To address this gap, we successfully introduced and characterized a novel cell stretching platform integrating tunable PDMS formulations using Sylgard 184 and 527 that mimic the physiological to pathophysiological range, allowing us to investigate combinatory effects of elastic properties and mechanical loading on AF cells. Other studies have reported similar approaches that covalently bonded polyacrylamide (PA) substrates with tunable stiffnesses on commercial stretching platforms from Flexcell and STREX [35, 41]. However, these approaches rely on the purchase of costly commercial stretching platforms or require a multi-step preparation process, including the fabrication of PDMS membranes, tuning the mechanical properties of PA, bonding PA to the PDMS surface, and subsequently modifying the surface chemistry to enable cell attachment. This process is time consuming, requires advanced surface chemistry using potentially hazardous chemicals, and the long-term mechanical stability of the PA-PDMS bond remains unclear. In contrast, our approach is simpler, requires fewer steps, minimizes the use of hazardous chemicals, and is therefore a more accessible and effective alternative.

In this study, Sylgard 184 and 527 formulations were prepared that successfully achieved a broad and distinct range of Young’s moduli. While literature reports variability in AF stiffness measurements, due to testing methods, anatomical location, and degeneration grade, our achieved range of elastic properties offers a representative spectrum for investigating cellular responses to stiffness ranges recapitulating a healthy or degenerative environment [17–19]. Similarly, previous research has demonstrated the tuneability of the elastic properties of Sylgard 527 and 184 formulations from 5 kPa (Sylgard 527 only) [32, 33] to 1.72 MPa (Sylgard 184 only) [32]. Although not tested in this study, others have described that varying the components A and B of Sylgard 527 is sufficient to tune mechanical properties. They presented a formula to calculate the resulting stiffness, leading in case of an equal ratio of A and B to ∼9 kPa [42], matching our Low stiffness formulation.

As soft tissue ECM exhibits viscoelastic properties, cells do not exclusively respond to elastic properties but also to viscous substrate components [43]. The AF is attributed to exhibit viscoelasticity, which enables the tissue to resist deformation during cyclic loading [44, 45] and influences matrix stress relaxation and energy dissipation, thereby impacting cytoskeletal tension, focal adhesion dynamics and differentiation behavior [46, 47]. Petet *et al.* [30] provided evidence that varying, the Sylgard 184 base to crosslinker ratio and Sylgard 184 to 527 ratio, allows partial independent tuning of G’ and G’’, but the achievable range is limited and a complete decoupling in PDMS remains challenging. Our DMA confirmed that the formulations significantly influence G’, G’’, and G*, with dominating elastic properties. This represents a limitation of our system highlighting the need for alternative formulations, or methods allowing independent control. Among formulations, the Low substrate exhibited the highest viscous component (tanδ = 0.28). Viscous contributions could in principle lead to creep, stress relaxation, residual strain, or reduced peak strains [46, 47], but these behaviors were not specifically tested in our study. Nevertheless, frequency sweeps demonstrated a linear viscoelastic response at 1 Hz, and the average surface strain across cycles in all three substrates confirmed reproducible and stable behavior, suggesting negligible viscous contributions in the system. The increased G* and elastic response in our High substrate (degenerative environment) resembles the observed shear behavior in tested AF tissue [48]. Furthermore, they reported that tanδ ranges between 0.1 and 0.7, aligning with our reported ranges. Although in degenerated AF samples no significant differences of the tanδ was observed compared to undegenerated tissue [48]. Overall, our findings share similarities with Petet *et al.* [30]. The authors reported a G’ of approximately 53 kPa for a comparable formulation to our Mid substrate, which is higher than our obtained G’ of 23.12 kPa, likely being impacted by discrepancies in curing conditions (110°C for 18 h vs. 65°C for 18 h). This highlights the challenges in comparing DMA data throughout literature.

While traditional Sylgard 184 is known to be biocompatible [26, 49], only few studies have suggested that Sylgard 527 formulations can equally promote cell adhesion and growth [30, 32, 34]. Our results confirm that all PDMS formulations support AF cell attachment at 24 h, cell growth over 72 h and exhibited negligible cytotoxicity. Although the High stiffness material supported cell growth, it was significantly lower than the Low stiffness material, indicating softer substrates may be favorable for cell growth. Furthermore, we excluded coating effects by comparing FN coating to COL. In addition to FN, COL is a widely used coating known to enhance cell adhesion and is physiological relevant due to the abundance of COL in the AF ECM [50, 51]. The different surface coatings (FN and COL) equally supported cell adhesion and growth, consistent with previous findings of similar cell proliferation of mesenchymal stem cells (MSCs) on plasma pre-treated FN and gelatin coated PDMS [28]. Other studies that compared various coatings on PDMS showed that FN was favorable for cell proliferation on PDMS [49] and also enhanced cellular attachment on polyurethan scaffolds [52], supporting its use as a primary coating in our study. Although, cell adhesion and growth was unaffected in our static conditions, literature reported that ECM based adsorption on PDMS may limit long-term cell growth and adhesion, due to relatively weak PDMS-ECM coating interaction being a potential result of the hydrophobic recovery effect [28, 53]. These studies suggest considering alternative surface functionalization methods for long-term studies such as polydopamine coatings [51], covalent ECM binding onto PDMS by using agents such as (3-Aminopropyl)triethoxysilane (APTES) with glutaraldehyde [54, 55] or Sulpho-Sanpah [56].

Aside from cell viability and attachment, analyzing cell spreading on PDMS is important as cell spreading has been shown with different cell types to tune cytoskeletal tension and integrin clustering activating mechanotransduction pathways such as RhoA/ROCK and YAP/TAZ. Consequently, changes in spreading area can remodel focal adhesion dynamics, shift gene expression programs, and influence behaviors like proliferation, differentiation, and migration [32, 34, 57, 58]. Although we observed significant differences in cell spreading, they are mild and varied between donors. However, it is consistent that the Low substrate generally promoted greater cell area, with opposite trends on High substrate, which is in accordance with the metabolic activity at 72 h. Stiffness-induced effects of PDMS on cell spreading reported in literature are conflicting. One study reported no distinctive differences between the cell spreading area of MC3T3-E1 pre-osteoblasts [34] cultured on Sylgard 184 and 527 formulations (with stiffness ranging from 0.6 MPa up to 2.7 MPa), yet stiffness dependent differences in vinculin levels, an indicator of cell-substrate interaction, were observed [59]. In contrast, others — using similar PDMS formulations — reported stiffness dependent effects on myotube length and clustering in rat adrenal pheochromocytoma cell lines [32]. Although these contradictory findings may reflect cell type specific behaviors, as well as variability in PDMS formulations or coatings, additional complex and interdependent factors are likely at play. For example, a study using adjusted Sylgard 184 substrates found that the presence of FBS in culture media minimized stiffness dependent effects on vascular smooth muscle cell spreading compared to serum-free conditions [60]. Although it has been speculated that Sylgard 184 and 527 may display differences in surface or chemical properties that could influence cell behavior, prior studies have clearly demonstrated that no differences in surface wettability, roughness, and energy after surface modification exist, resulting in similar protein adsorption on the surfaces. These findings also suggest that porosity is not altered which we confirmed through SEM imaging (data not shown), suggesting no detectable signs of porosity at the surface level. These studies also report similar chemical structures between the formulations, with assumed differences primarily in the polymer chain length, which supports our findings of nearly equal chemical profiles in FTIR-ATR spectra [30, 32].

Maintaining cell viability and adhesion is crucial, yet challenging during load applications, especially when trying to simulate detrimental (i.e. excessive or prolonged) loading, a critical factor in IVD pathology, contributing cellular stress, inflammation, and ultimately reducing cellular viability [14, 61]. Our findings indicate that mechanically stimulated AF cells remain viable for up to 14 h of loading, with a marked decline by 18 h, suggesting a threshold beyond which mechanical stress impairs survival. While the precise cause of the reduced viability was not examined, the strength of cell adhesion to the PDMS substrate may be a contributing factor.

To our knowledge, this study is the first to comprehensively characterize STREX stretching chambers. Tensile testing confirmed that the stiffness of these chambers exceeds the physiological and pathophysiological stiffness range reported in the AF [17–19, 48]. Contrary to assumptions of uniform strains along the loading axis on stretching chambers, DIC revealed two strain components at the membrane surface: an axial strain along the loading direction and a transverse strain acting perpendicular to it accompanied with non-uniform strain distributions across the stretching chamber surface. Notably, at 20% applied deformation, only about 60% of the total strain, as shown in Figure 6C, was attributed to the axial component, highlighting a significant transverse strain contribution. These findings point out that depending on where the cells are located, they experience different strain patterns varying in both magnitude and type of strain. The observed strain pattern stems from PDMS’s near-incompressibility (Poisson’s ratio ≈0.5 [62]), whereby axial tension produces substantial lateral contraction. These findings align with a previous finite element analysis that reported similar transverse strain components during PDMS membrane stretching [63]. The observed non-uniform strains in the commercial and custom chambers were most pronounced in the Low stiffness, because it has reduced resistance to deformation therefore producing higher surface strains accompanied by larger strain heterogeneity within the chamber. Nevertheless, experimental repeatability is not compromised, as average strain patterns at extreme magnitudes of 20% remained consistent between chambers. Our measurements revealed that the actual strain experienced by cells deviated from the applied ε*_eng_* and is dependent on the chamber type, making direct comparison among studies challenging. These findings highlight the importance of carefully characterizing surface strains in self-fabricated chambers, as both chamber geometry and stiffness can influence strain patterns and magnitude. Such characterization raises the awareness of the present strain patterns and informs experimental design and interpretation of results in future studies. Finally, our approach and data provide a foundation for improved designs, including the use of finite element simulations and experimental testing. As this study focused on single, short-term (up to 18 h) mechanical testing, a systematic evaluation of the substrate’s mechanical stability and fatigue was not performed because no monitored visual signs of mechanical failure such as surface damage or chamber leakage was observed. This is supported by Bernardi *et al*. [64], who reported negligible stress decay over 200,000 cyclic loadings of Sylgard 184. Consequently, Low PDMS is not deemed to exhibit higher fatigue risk, as it experiences lower internal stresses under applied strain compared to the High substrate. Wang *et al.* [65] confirmed low hysteresis in soft and hard PDMS (Sylgard 184), while soft substrates delayed rupture risk. Furthermore, the 20% strain applied is well below the expected rupture strain of the PDMS formulations (>100%) [32]. Additionally, in case of present surface cracks, softer substrate tends to redistribute the stress around the cracks more efficiently, producing blunt crack tips, hence reducing failure risk. Although our results suggest promising mechanical performance, future studies should systematically test mechanical stability and fatigue resistance of the formulations, particularly in extended and more extreme loading conditions.

To explore how the strain of our custom chambers affects AF cell behavior, we examined cellular orientation under cyclic stretching. Cellular reorientation minimizes the impact of external stress and restores tensional homeostasis, a concept regulated by dynamic remodeling of stress fibers and focal adhesions mediated by Rho GTPase and myosin light chain [35, 66, 67]. Strain application of 5.8% measured strain (ε*_eng_* = 8%, 1 Hz) on High chamber was sufficient to induce cell alignment within 6 h. This is consistent with previous studies reporting initial cell alignment after 3 h of mechanical loading at 15% strain and 2 Hz [63] and another study suggested that alignment is sensitive to the strain frequency, amplitude, and duration with a steady-state alignment reached at ∼4 h (8% and 2 Hz) [68]. Quinlan *et al*. [35] reported alignment of vascular interstitial cells on stiff (50 kPa) PA substrates, but alignment was attenuated on very soft (0.3 kPa) PA substrates due to reduced spreading and traction forces. In contrast, our bovine AF cells spread well on the Low substrate (10 kPa), indicating sufficient actomyosin contractility and FA turnover, resulting in adequate prestress and adhesion strength required for cellular reorientation. This supports the conclusion that perpendicular alignment is primarily strain dependent. Nevertheless, alignment on the Low substrate was significantly reduced compared to the High stiffness chambers, likely due to weaker force transmission from the substrate to the cells. This process is relevant in the AF, where structural integrity and susceptibility to loading is dependent on the highly aligned collagen I fibers oriented perpendicularly to the experienced tensile loading [13]. Such a behavior may represent an adaptive response of the AF cells to cyclic loading, reducing sensitivity to high strains while maintaining alignment as mechanical loading promotes matrix deposition. Interestingly, we have shown that alignment was more pronounced in regions experiencing higher degree of axial to transverse strain distribution, while areas with higher relative transverse strain contributions showed reduced alignment. These findings may suggest that cells are highly sensitive to the complex strain patterns such as magnitude and directionality of the experienced strain, as revealed by DIC, but further investigations are needed.

Collectively, this study presents a novel PDMS based stretching platform by integrating a range of elastic properties relevant to physiological and pathophysiological conditions, with potential applications beyond the IVD. We demonstrate the feasibility of the system and provide a detailed analysis of strain distribution in both custom-fabricated and commercial STREX chambers. While these systems effectively transmit strain to the AF cells, an in-depth biological evaluation at the cellular level was beyond the scope of this study. Future work will, therefore, focus to understand how strain patterns and substrate stiffness affects AF-cell responses at the gene and protein expression level, as well as evaluating alternative surface treatments (e.g., Sulfo-Sanpah-based ECM coating).

## 5 Declarations

### 5.1 Author contributions

Johannes Hasler conceived and designed the study, conducted experiments, analyzed data, and drafted the manuscript. Mikkael Lamoca conceptualized the design of the stretching chambers, performed tensile testing and helped with figures. Christopher L. Lewis and Kory Schimmelpfennig provided input for material testing and supported DMA and FTIR-ATR testing of the PDMS substrates. Shuhuan Zhang contributed with supervision and extensive training of chamber characterization using digital image correlation. Rui Liu contributed with the conception of strain characterization and provided Lavision’s StrainMaster equipment. Wolfgang Hitzl conducted the statistical analysis. Sami Farajollahi contributed to the design, implementation and execution of the experimental method for the cellular alignment investigation. Vinay V. Abhyankar supervised the study and provided critical scientific input on the design, fabrication and application of the PDMS based system as well as contributing to the interpretation of results. Karin Wuertz-Kozak originated the research concept, provided funding and project management, and supported data interpretation. All authors contributed to revising the manuscript and approved the final version.

## 5.3 Acknowledgement

This work was supported by the National Institutes of Health: 5R16GM146717 (Johannes Hasler, Mikkael Lamoca and Karin Wuertz-Kozak), R16GM146687 (Sami Farajollahi and Vinay V. Abhyankar), U2CAG088071 (Sami Farajollahi and Vinay V. Abhyankar), and the National Science Foundation: CBET 2150798 (Vinay V. Abhyankar). The authors thank Gabbie Wagner for the support with bovine AF cell isolation.

## 5.4 Availability of raw data and materials

The raw data and material, generated in this study, are publicly available in the Figshare database: https://doi.org/10.6084/m9.figshare.28883804

