## Supplementary Material for "Development and Characterization of a Tunable PDMS Substrate Model for Investigating Elastic Properties and Mechanical Stretching in Intervertebral Disc Cells"

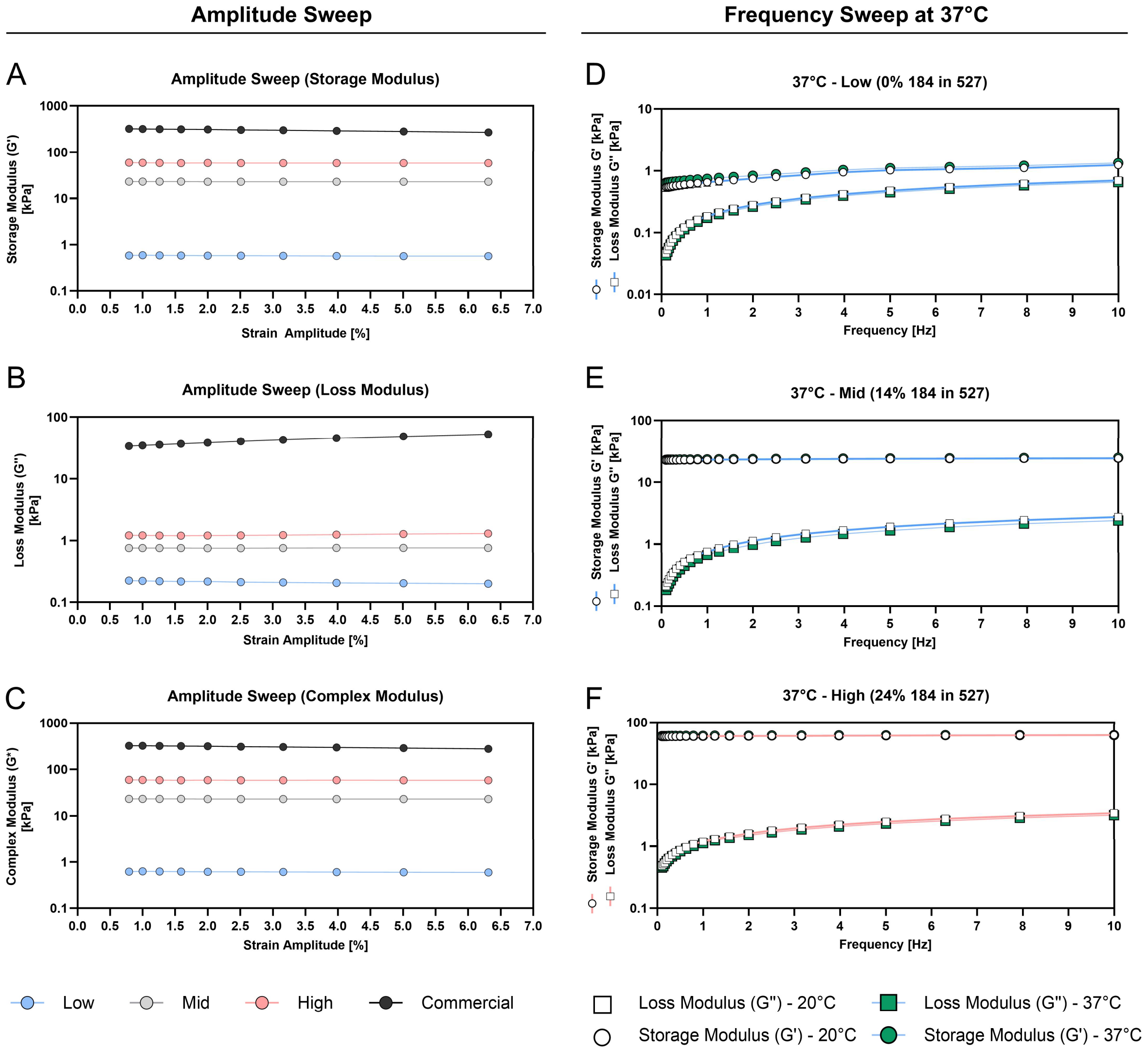


**Supplementary Figure 1:** Dynamic mechanical assessment (DMA) of the PDMS formulations prepared with varying ratios of Sylgard 184 to Sylgard 527 (0, 14, 24 or 100% Sylgard 184). **A-C)** Amplitude sweep (0.79% – 6.31%) in the linear viscoelastic region, including storage modulus **(A)**, loss modulus **(B)**, and complex modulus **(C)** for PDMS formulations containing 0%, 14%, 24%, and 100% Sylgard 184 in Sylgard 527 (*n* = 1). **D-F)** Frequency sweep (0 – 10 Hz) at 20 °C, comparing storage and loss moduli for PDMS formulations: Low **(D)**, Mid **(E)**, and High **(F)** at 37 °C (*n* = 1).

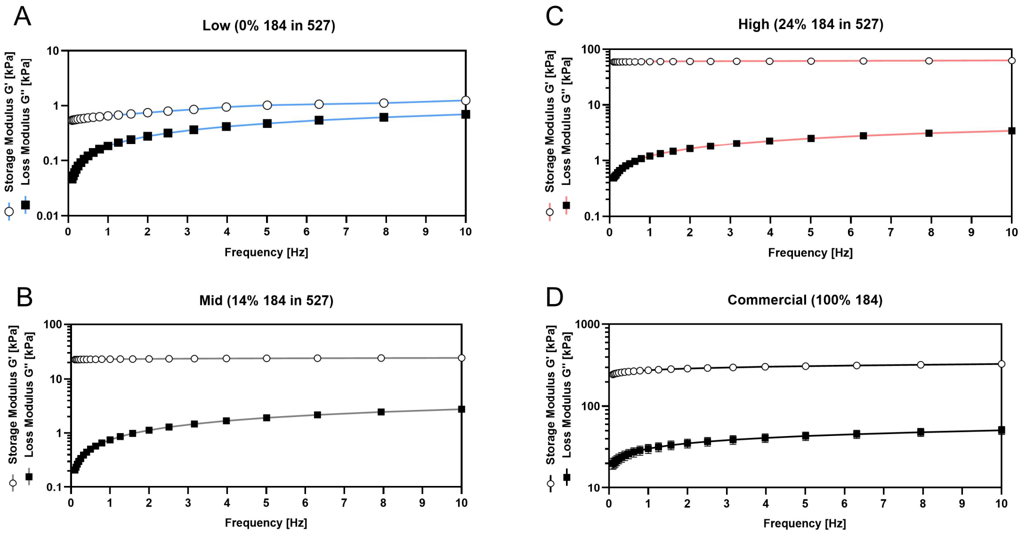


**Supplementary Figure 2:** Dynamic mechanical assessment (DMA) of the PDMS formulations prepared with varying ratios of Sylgard 184 to Sylgard 527 (0, 14, 24 or 100% Sylgard 184). **A-D)** A frequency sweep between frequencies of 0 Hz – 10 Hz was performed to assess storage- and loss modulus of the PDMS formulations: Low **(A)**, Mid **(B)**, High **(C)**, and Commercial **(D).** (*n* = 3), mean ± SD.

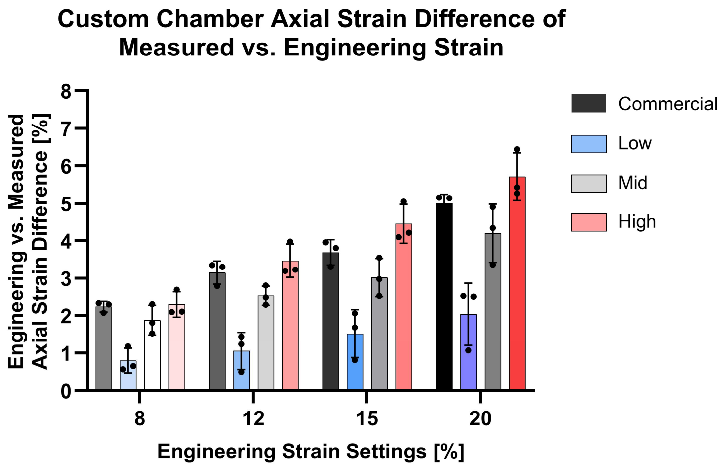


**Supplementary Figure 3:** Differences of measured strain and engineering strain settings (8-20%) in percentages. (*n* = 3), mean ± SD.

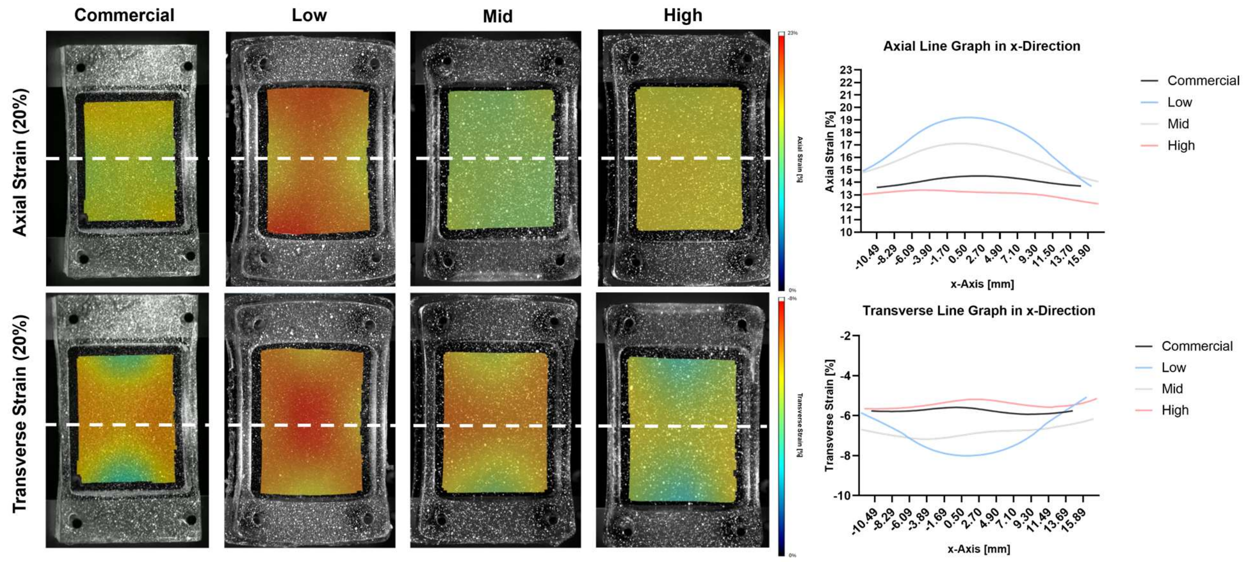


**Supplementary Figure 4:** Axial (top row) and transverse (bottom row) strain distributions at 20% applied strain ($\varepsilon_{eng}$) of representative chambers (Commercial, Low, Mid and High). White dashed line indicates the location used for the line graphs for axial and transverse strain along the x-Axis (mid of y-Axis).

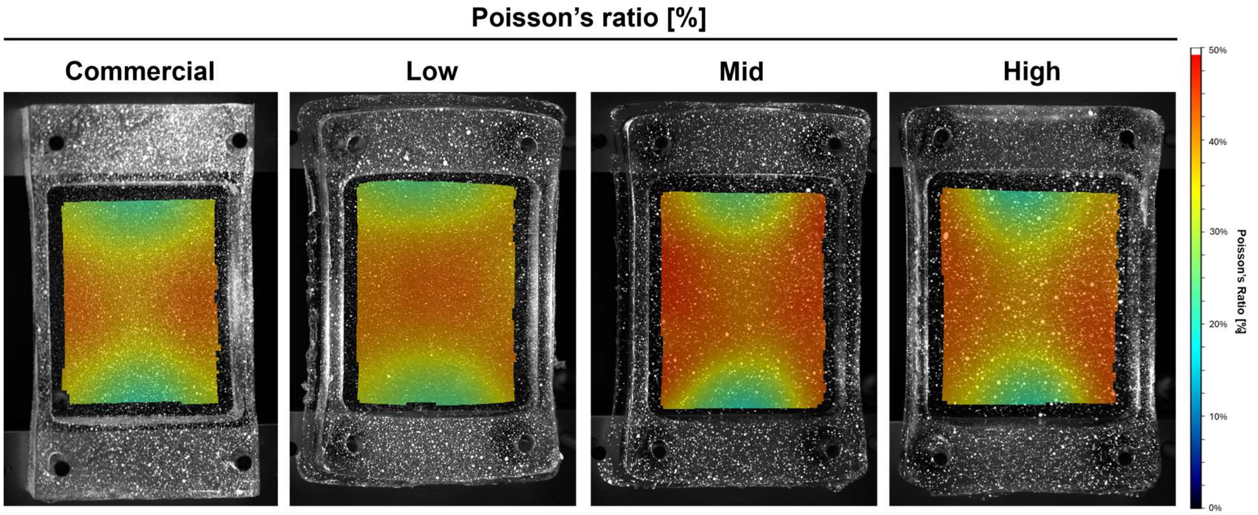


**Supplementary Figure 5:** Poisson's ratio distribution (%) on the cell interface surface at 20% $\varepsilon_{eng}$, illustrating relative transverse and axial strain contributions. Higher percentages highlight larger contribution of transverse strain compared to axial strain. (*representative images* of Commercial, Low, Mid and High chambers).

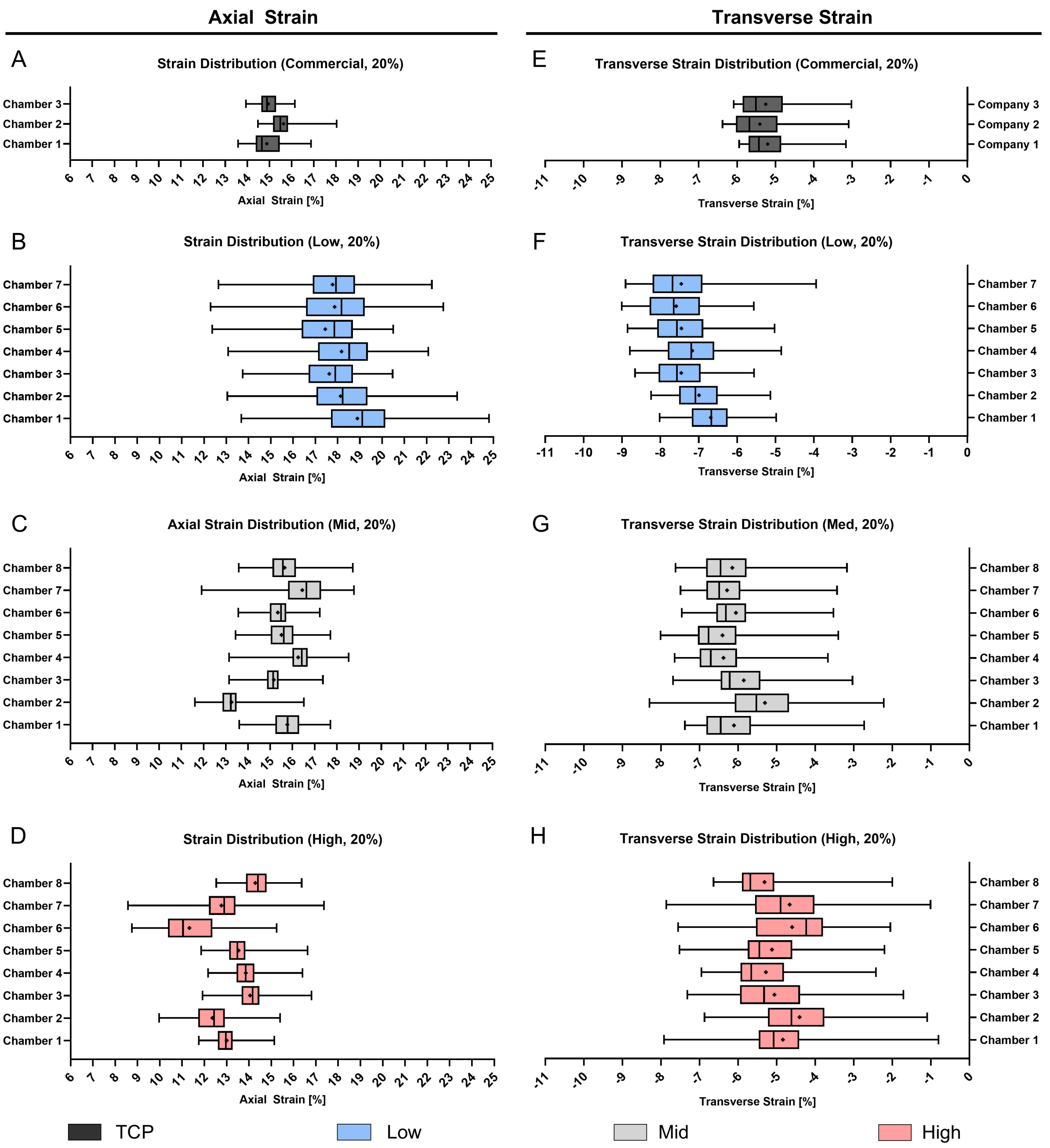


**Supplementary Figure 6:** Frequency distribution plot of individual strain distribution characterization within a stretching chamber at maximum engineering strain settings of 20% represented as box plots. **A-D)** Axial strain distribution within the cell interaction surface of the PDMS chambers for commercial **(A)** and custom chambers **(B-D)**. **E-H)** Axial strain distribution within the cell interaction surface of the PDMS chambers for **E)** commercial and **F-H)** custom chambers.

**Supplementary Table 4:** Descriptive statistics of the axial and transverse strain distribution in the stretching chambers including Low, Mid, High and commercial chambers. Data includes the strain range and standard deviation (SD) as well as an overall average of both in percentage.

| | Commercial | | Low | | Mid | | High | |
|---|---|---|---|---|---|---|---|---|
| | **Axial** | | | | | | | |
| | **Range [%]** | **± SD [%]** | **Range [%]** | **± SD [%]** | **Range [%]** | **± SD [%]** | **Range [%]** | **± SD [%]** |
| **Chamber 1** | 3.30 | 0.73 | 11.12 | 1.92 | 4.11 | 0.75 | 3.38 | 0.61 |
| **Chamber 2** | 3.55 | 0.78 | 10.32 | 1.84 | 4.91 | 0.75 | 5.42 | 1.02 |
| **Chamber 3** | 2.21 | 0.45 | 6.74 | 1.48 | 4.22 | 0.68 | 4.89 | 0.81 |
| **Chamber 4** | - | - | 8.99 | 1.81 | 5.39 | 0.82 | 4.24 | 0.70 |
| **Chamber 5** | - | - | 8.12 | 1.76 | 4.27 | 0.75 | 4.77 | 0.73 |
| **Chamber 6** | - | - | 10.46 | 2.12 | 3.67 | 0.63 | 6.48 | 1.32 |
| **Chamber 7** | - | - | 9.59 | 1.61 | 6.86 | 1.32 | 8.80 | 1.29 |
| **Chamber 8** | - | - | - | - | 5.13 | 0.83 | 3.83 | 0.68 |
| **Mean** | **3.02** | **0.65** | **9.33** | **1.79** | **4.82** | **0.82** | **5.23** | **0.89** |
| | **Transverse** | | | | | | | |
| **Chamber 1** | 2.78 | 0.64 | 3.03 | 0.69 | 4.65 | 0.99 | 7.13 | 0.94 |
| **Chamber 2** | 3.29 | 0.78 | 3.11 | 0.71 | 6.08 | 1.17 | 5.78 | 1.15 |
| **Chamber 3** | 3.07 | 0.74 | 3.10 | 0.74 | 4.66 | 0.96 | 5.60 | 1.17 |
| **Chamber 4** | - | - | 3.95 | 0.86 | 3.97 | 0.88 | 4.53 | 0.96 |
| **Chamber 5** | - | - | 3.83 | 0.84 | 4.61 | 0.92 | 5.32 | 1.02 |
| **Chamber 6** | - | - | 3.44 | 0.85 | 3.94 | 0.75 | 5.50 | 1.16 |
| **Chamber 7** | - | - | 4.97 | 0.95 | 4.06 | 0.78 | 6.86 | 1.31 |
| **Chamber 8** | - | - | - | - | 4.44 | 0.94 | 4.64 | 0.98 |
| **Mean** | **3.05** | **0.73** | **3.63** | **0.81** | **4.55** | **0.92** | **5.67** | **1.09** |

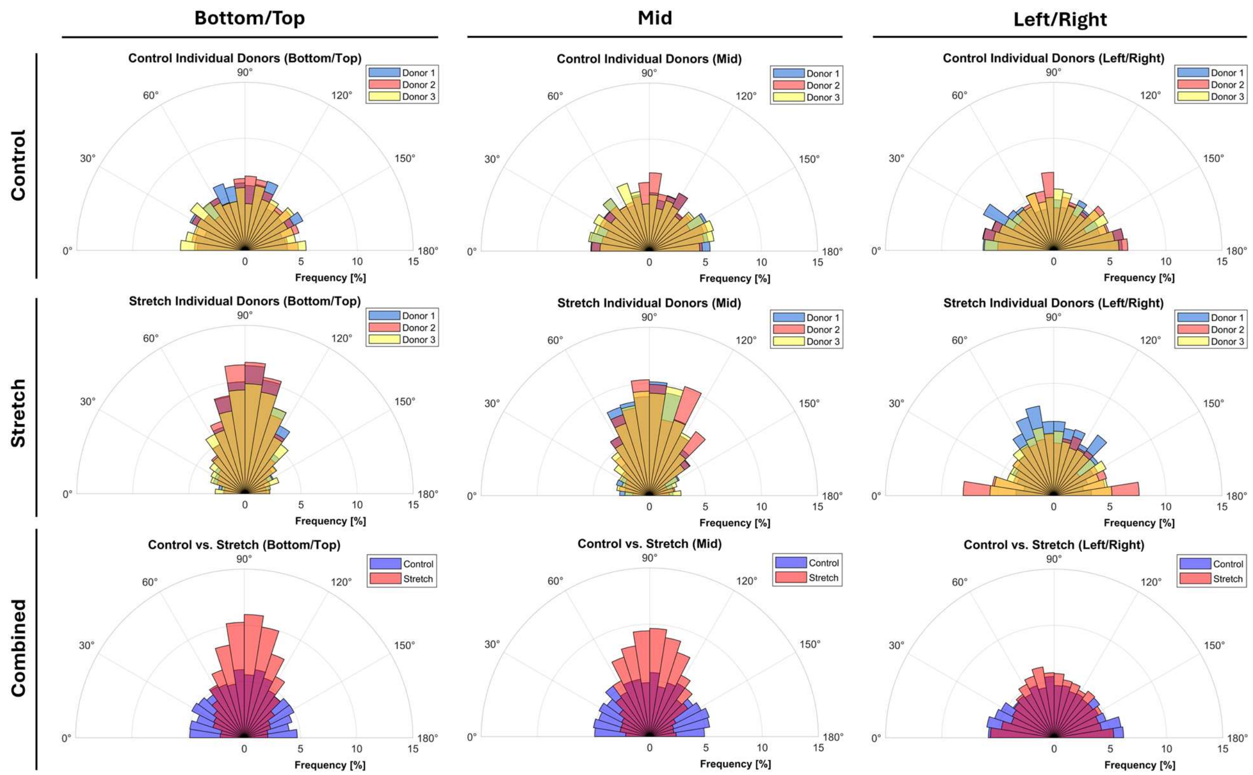


**Supplementary Figure 7:** Cellular alignment under cyclic stretching in High stiffness chambers. Different locations within the stretching chamber were imaged including high strain location (Top/Bottom, Mid) and low strain locations (Left/Right). Data is presented as an overlay polar histogram separately for three individual Donors (Control and Stretch) and as average plots combining control, stretch and Donors in one figure (combined) for three donors and three images at different locations. A polar histogram presents the cellular alignment of unstretched (control) and stretched chambers.
